# Small Molecule Antiviral Compound Collection (SMACC): a database to support the discovery of broad-spectrum antiviral drug molecules

**DOI:** 10.1101/2022.07.09.499397

**Authors:** Holli-Joi Martin, Cleber C. Melo-Filho, Daniel Korn, Richard T. Eastman, Ganesha Rai, Anton Simeonov, Alexey V. Zakharov, Eugene Muratov, Alexander Tropsha

## Abstract

Diseases caused by new viruses costs thousands if not millions of human lives and trillions of dollars in damage to the global economy. Despite the rapid development of vaccines for SARS-CoV-2, the lack of small molecule antiviral drugs that work against multiple viral families (broad-spectrum antivirals; BSAs) has left the entire world’s human population vulnerable to the infection between the beginning of the outbreak and the widespread availability of vaccines. Developing BSAs is an attractive, yet challenging, approach that could prevent the next, inevitable, viral outbreak from becoming a global catastrophe. To explore whether historical medicinal chemistry efforts suggest the possibility of discovering novel BSAs, we (i) identified, collected, curated, and integrated all chemical bioactivity data available in ChEMBL for molecules tested in respective assays for 13 emerging viruses that, based on published literature, hold the greatest potential threat to global human health; (ii) identified and solved the challenges related to data annotation accuracy including assay description ambiguity, missing cell or target information, and incorrect BioAssay Ontology (BAO) annotations; (iii) developed a highly curated and thoroughly annotated database of compounds tested in both phenotypic (21,392 entries) and target-based (11,123 entries) assays for these viruses; and (iv) identified a subset of compounds showing BSA activity. For the latter task, we eliminated inconclusive and annotated duplicative entries by checking the concordance between multiple assay results and identified eight compounds active against 3-4 viruses from the phenotypic data, 16 compounds active against two viruses from the target-based data, and 35 compounds active in at least one phenotypic and one target-based assay. The pilot version of our SMACC (Small Molecule Antiviral Compound Collection) database contains over 32,500 entries for 13 viruses. Our analysis indicates that previous research yielded very small number of BSA compounds. We posit that focused and coordinated efforts strategically targeting the discovery of such agents must be established and maintained going forward. The SMACC database publicly available at https://smacc.mml.unc.edu may serve as a reference for virologists and medicinal chemists working on the development of novel BSA agents in preparation for future viral outbreaks.

## Introduction

Infectious diseases have had profound impacts on global human health since the beginning of time. In the past two decades factors such as population growth and travel have increased the rate of viral outbreaks, with a new viral threat seen nearly every year.^1^ This includes the emergence of severe acute respiratory syndrome coronavirus (SARS-CoV), Middle East respiratory syndrome (MERS-CoV), Zika virus disease, Ebola virus disease, and variety of influenza strains (H5N1, H7N9, H1N1, etc.). These viruses are just a handful of over 200 viral species annotated by the International Committee for Taxonomy of Viruses as threats to human health.^2^ The millions of lives lost and trillions of dollars in damage to the global economy due to the recent pandemic caused by severe acute respiratory syndrome coronavirus 2 (SARS-CoV-2) highlight the importance of medications that can offer protection against diverse viral threats that can emerge in the future.^3^

It is evident from the current SARS-CoV-2 outbreak that scientific advances have increased our capabilities for rapid development of new vaccines. However, none of the vaccines developed thus far offers 100% protection. Furthermore, viral evolution poses a threat to further decrease the vaccines efficacy, additionally, widespread hesitancy against vaccination leaves a large majority of the population susceptible to the viral disease.

Broad-spectrum antiviral (BSA) drugs could protect against emergent viruses; however, the development of such drugs has been challenging. The standard drug development and clinical testing process averages 10–15 years; thus, it is impossible to see an immediate emergence of new, effective drugs following an outbreak. As of today, there are only 90 approved antiviral drugs of which 11 are approved to treat more than one virus.^4^ One could speculate that the ability of the 11 drugs to be effective against more than one virus could be explained by the conservation of their targets or mechanisms of action. For example, acyclovir triphosphate competes with dGTP to inhibit viral DNA polymerase activity in two human neurotropic alpha herpesviruses, herpes simplex virus and varicella zoster virus.^4^ Most approved antiviral drugs are effective against herpes, hepatitis, or human immunodeficiency viruses, but offer no protection against the recent SARS-CoV-2 pandemic. However, the fact that medications active against more than one virus do exist fuels the hypothesis that such medications can be developed in principle via concerted strategic effort.

Despite the clear need for BSA medications, previous outbreaks have shown that the interest in supporting viral research and drug discovery vanishes quickly after about a year past the viral threat, leaving the work toward an effective medication unfinished.^5^ A good example is the history of Paxlovid, a recently approved Pfizer medication against SARS-CoV-2. The respective drug candidate was initially discovered to work against SARS in 2002-2003 by inhibiting the virus’ main protease (3CL-Pro) but its further development was frozen after SARS vanished. When SARS-CoV-2 emerged, and it was quickly discovered that its main protease, especially its active site, is almost identical to its counterpart in the original SARS, the initial drug development program was restarted and Paxlovid was relatively quickly developed by Pfizer through focused medicinal chemistry optimization efforts.^6^ This story clearly indicates that there is a strong need for ongoing and well-funded research programs focused on the rational discovery of BSA drugs.

More than 380 trillion different viruses exist inside the human virome, but so far only about 200 have been considered harmful for human health.^7^ To support focused development of BSAs and learn from history, in this study we endeavored to collect, curate, and integrate all publicly accessible data on compounds tested in both phenotypic and target-based assays for emerging viruses of concern. To this end, we have (i) conducted a comprehensive evaluation of viruses holding the greatest potential threat to global human health, (ii) used the data available in ChEMBL, an online collection of bioactive molecules with drug-like properties, to build a curated, annotated, and publicly available database of compounds tested in both phenotypic and target-based assays for these viruses, and (iii) identified the most promising candidates with potential BSA activity. We dubbed this database Small Molecule Antiviral Compound Collection (SMACC) and made it publicly available online at https://smacc.mml.unc.edu. We expect that SMACC database can support further computational and experimental medicinal chemistry studies targeting rational design and discovery of novel BSAs.

## Methods

### Selection of viruses of interest and initial database generation

The ability of an infectious disease pathogen to cause a pandemic is impacted by several intrinsic characteristics, including the mode and timing of transmission, host population susceptibility, lack of effective therapeutic interventions or control measures, among others. Microbial pathogens infect humans through many routes of transmission, including through animal vector, fecal-oral or respiratory. Respiratory transmitted diseases are more likely to possess pandemic potential, as interventions to block human-to-human transmission via aerosols are more challenging to implement. The timing of disease transmission also impacts the spread of a disease, if a pathogen is transmissible early in the course of disease, especially if an individual is asymptomatic, this greatly facilitates potential for spread.

Viruses with a high replication rate, especially coupled with mutability of RNA and segmented RNA can rapidly gain attributes, including increased transmission or evading preexisting immunity, which also facilitates outbreak or pandemic spread. Viruses with high pandemic potential include *Coronaviridae, Paramyxoviridae*, *Bunyavirales*, *Picornaviridae*, *Filoviridae*, *Togaviridae,* and *Flaviviridae* virus families. Thus, we selected the following 13 viruses representative of five families to query respective chemical bioactivity data in ChEMBL: *Coronaviridae* (SARS-CoV-2, MERS-CoV, HCoV-229E), *Orthomyxoviridae* (H1N2, H7N7), *Paramyxoviridae* (RSV, HPIV-3), *Phenuiviridae* (Sandfly Fever), and *Flaviviridae* (Dengue, Zika, Yellow Fever, Powassan, West Nile).

All data was extracted from ChEMBL 29.^8^ The virus name, and any known alias were used as keywords to extract all phenotypic and target-based assays for each virus. For the target-based assays, we ran an additional search using virus and target name as the keywords to ensure no respective viral data was lost. To identify drug targets for each virus we searched existing literature using the keywords “ [virus_name] virus drug targets.” After extraction, the data for each virus were pre-processed and curated as described below. When examining the resulting datasets, we have identified a need for additional curation of assay annotations as discussed in the Results section.

### Data Curation

We followed protocols for chemical and biological data curation described by Fourches et al.^9–11^ In brief, specific chemotypes were normalized. Inorganic salts, organometallic compounds, and mixtures were removed. Duplicate compound entries were kept to define the activity calls for compounds tested against the same virus but using different assay protocols. Additionally, keeping duplicates in the database can be important for analyzing the overlap of compound activity between different viruses in a search for potential BSAs. However, mindful of computational modeling studies that require the removal of duplicate compound entries, these entries were carefully annotated in our database. All steps of data curation and integration were performed in KNIME v.4.1.4^12^ integrated with python v.3.7.3, RDKit v.4.2.0, and ChemAxon Standardizer v. 20.9 (ChemAxon, Budapest, Hungary, http://www.chemaxon. com). We summarized our database entries after curation in **Table S1**.

### Identification of compounds with multiple antiviral activity

A threshold of 10 µM, irrespective of the type of activity measurement, was applied to define the outcome (i.e., if a compound was active or inactive). When compound activity was reported with ambiguous operators (greater than, “>”, or less then, “<”, certain value), it was annotated as inconclusive. The final definition of the activity call for each compound was based on the concordance of all compound replicate entries tested against the same virus but in different assays (or the same viral target for the target-based dataset). Three outcomes were possible: (i) the compound was active when tested in all assays; (ii) it was active in some assays and inactive in others; and (iii) it was inactive in all assays. In case (i), the compound was considered active while in case (iii), inactive. Any compound in case (ii), with discordant activity calls resulting from different assays, i.e., with at least one activity call different from other ones for the same virus, was considered inconclusive and was not used for the overlap analysis to identify compounds with multiple antiviral activity. A compound was also annotated as inconclusive if the assay reported the compounds activity as “Not Determined.” Finally, all compounds tested in different viruses (or viral targets in the target-based dataset) were analyzed and those showing activity against two or more viruses were selected as potential BSAs. Table 1 summarizes our protocols to decide on the final activity calls for compounds included in SMACC database.

**Table 1.** Rules for making the final activity calls for compounds in SMACC database.

| Virus | Assay | Cell Type | Activity discordance* | Standard Value | Activity Call |
| --- | --- | --- | --- | --- | --- |
| same | same | same | no | <10 uM or >50% | active |
| same | same | same | no | >10 uM or <50% | inactive |
| any | any | any | no | “Not Determined” | inconclusive |
| same | same | same | yes | Any | inconclusive |
| same | different | same | yes | Any | inconclusive |
| same | different | different | yes | Any | inconclusive |

\* At least one discordant duplicate/replicate compound

### Cluster Analysis

The curated compound structures were submitted to hierarchical cluster analysis in KNIME v.4.1.4^12^ integrated with python v.3.7.3 (SciPy and Matplotlib libraries) and using RDKit descriptors (RDKit v.4.2.0). The optimal number of clusters was determined by the software default Euclidean distance cut-off.

## Results

### Ontological examination and curation of assays reported in ChEMBL

#### Phenotypic assays

While ChEMBL does an exceptional job at providing the largest curated and publicly available bioactivity database, we have identified multiple issues requiring additional data curation efforts to yield a clean database of antiviral activity data. Specifically, we found that phenotypic assays for antiviral compounds have been annotated in ChEMBL with inconsistent ontological annotation, which creates uncertainty in the data. The most common finding was the inconsistent use of BioAssay Ontology annotations for the assay type. As stated on the BioAssay Ontology’s homepage,^13^ “The BioAssay Ontology (BAO) describes chemical biology screening assays and their results including high-throughput screening (HTS) data for the purpose of categorizing assays and data analysis.” In practice, proper BAO usage has been considered a universal best practice and highly trusted by users. However, in the phenotypic assay data collection from ChEMBL, the misuse of the BAO ontology was evident: for 9 of 13 viruses the assay type was recorded in ChEMBL as “Organism-Based” rather than “Cell-Based”. This was concerning because a virus does not meet the criteria of a living organism as its life cycle relies on the host organism. Therefore, these assays should be properly reported as cell-based; thus, we corrected their annotation respectively in our database. The impact of this round of curation on the quality and usability of the extracted data was dramatic. Indeed, in the absence of such manual analysis and correction of mis-annotated data, if one were to search ChEMBL for “cell-based assays” for these viruses, 99.44% (27,410 of 27,562 entries) of the data would have been uncovered. This analysis indicates a critical importance of careful data processing by chemical bioactivity data curators for both the accuracy of chemical structures (which has substantially improved over the years^14,15^) and correctness of activity labeling such that users can obtain the entirety of <u>existing</u> but effectively, <u>hidden</u> data for which they searched.

Missing, i.e., absent from their designated entry field, data annotations were also extremely common. For example, despite there being a distinct field for the respective entry, 13.72% of all phenotypic assays results did not indicate which cell type was used. Instead, we found the records of the cell type in the assay descriptions, which allowed us, in this case, to extract and properly annotate this field. However, 36.73% of all missing cell types were not listed in the assay description either, leaving one searching for the exact assay in the linked paper and trying to identify the cell type used, which is what we had to do. This process had to be done manually, which made it extremely time consuming, and, in some cases, no clear cell type could be identified. If the cell type was not identified eventually, it was annotated as “unclear.” These cases are reported in **Table 2** as “Cell Type Completely Missing”. Yet, this tedious work resulted in the additional recovery of ∼4% of all phenotypic assay results. Another issue of missing data annotations was uncovered when we looked into the class of assays. Most assays were not labeled to indicate whether they were primary, counter, or cytotoxicity assays. Furthermore, the assay descriptions also failed to provide an appropriate level of detail. Many assay descriptions simply reported “Antiviral activity against virus X.” Such descriptions are missing information on assay conditions like time, substrate, equipment as well as cell type and purpose of the assay. Lacking such details makes it impossible to analyze data reproducibility and prohibits meaningful integration of multiple assay results. **Table 2** summarizes of our effort to procure and enrich the original annotation of data found in ChEMBL.

**Table 2.** Curation issues of phenotypic data

| Virus | # of Entries | # of Assays | Incorrect BAO Assay Annotation | Assay Type Missing in Description | Cell Type Missing | Cell Type Available in Description | Cell Type Completely Missing |
| --- | --- | --- | --- | --- | --- | --- | --- |
| SARS-CoV-2 | 18,190 | 21 | 18,190 | 2 | 7 | 0 | 7 |
| MERS-CoV | 49 | 9 | 49 | 0 | 49 | 49 | 0 |
| HCoV-229E | 164 | 11 | 164 | 7 | 69 | 65 | 5 |
| Dengue | 2,685 | 581 | 2,685 | 191 | 1,682 | 1,495 | 187 |
| Yellow Fever | 930 | 66 | 930 | 41 | 169 | 72 | 97 |
| Zika | 357 | 91 | 357 | 19 | 334 | 308 | 26 |
| West Nile | 514 | 102 | 514 | 81 | 308 | 148 | 160 |
| Powassan | 24 | 1 | 0 | 0 | 0 | 0 | 0 |
| RSV | 2,906 | 239 | 2,862 | 586 | 608 | 141 | 467 |
| HPIV-3 | 1,632 | 161 | 1,566 | 435 | 520 | 96 | 421 |
| H1N2 | 61 | 7 | 43 | 0 | 18 | 18 | 0 |
| H7N7 | 26 | 7 | 26 | 0 | 18 | 0 | 18 |
| Sandfly Fever | 24 | 3 | 24 | 0 | 2 | 0 | 2 |

#### Target based assays

While many of the issues discussed above for phenotypic assays were not present in the target-based assays, there were some cases that needed further attention. One example includes 536 entries deposited as compounds tested against “genome polyprotein” of West Nile or Zika viruses. However, upon closer examination of the ChEMBL records, we have established that these compounds were actually tested against the NS2B-NS3 Protease, rather than the entire genome polyprotein.

To summarize this section, when using ChEMBL as a curated^16^ source of data on antiviral compounds, we have uncovered multiple special issues with inconsistency or mislabeling (cf. Table 1) of the biological assays data. We have addressed these issues by assigning correct BAO annotation to the data extracted from ChEMBL to enable the creation of a refined specialized SMACC database of antiviral compounds tested in diverse antiviral assays.

### Development of the curated data entries in the SMACC database

Our extensive curation efforts following the protocols described in Methods led to the removal of compounds from each viral family in the phenotypic data set (**Figure 1**). From the initial compound list through the normalization of specific chemotypes, we removed the ∼26 % of compounds (n=4801) from *Coronaviridae* family, 25% (n=6) from the *Phenuiviridae*, ∼21% (n=594) from *Flaviviridae* and ∼8% (n=7) from *Orthomyxoviridae* and ∼8% (n=350) from *Paramyxoviridae*.

**Figure 1.**
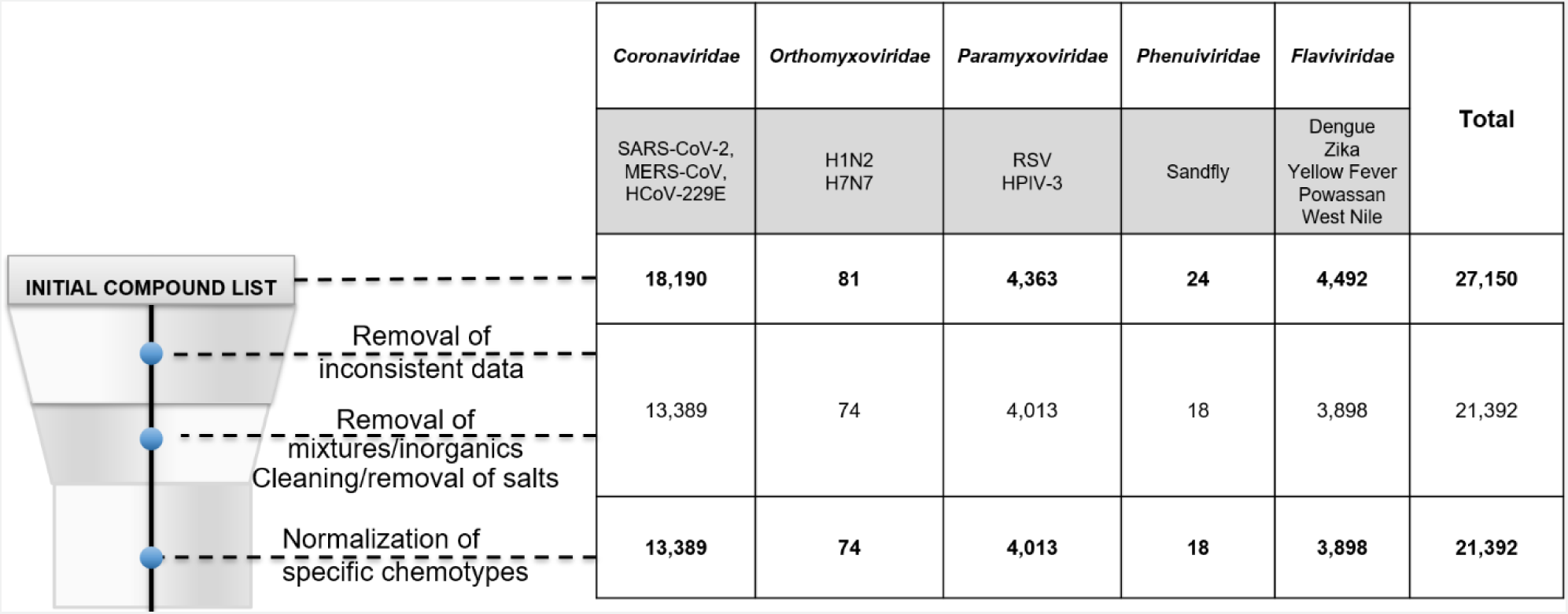
The effect of phenotypic assay data curation on reducing the resulting dataset sizes.

For the target-based curation, there were less compounds removed due to inconsistent data and molecular cleaning (**Figure 2**). Here the family where the largest number of compounds were removed was also the *Coronaviridae* which was less than 1% of all compounds (n=13). Interestingly, our annotation efforts followed a similar pattern, where the target-based data were deposited with more annotations, therefore requiring less curation than the phenotypic data.

**Figure 2.**
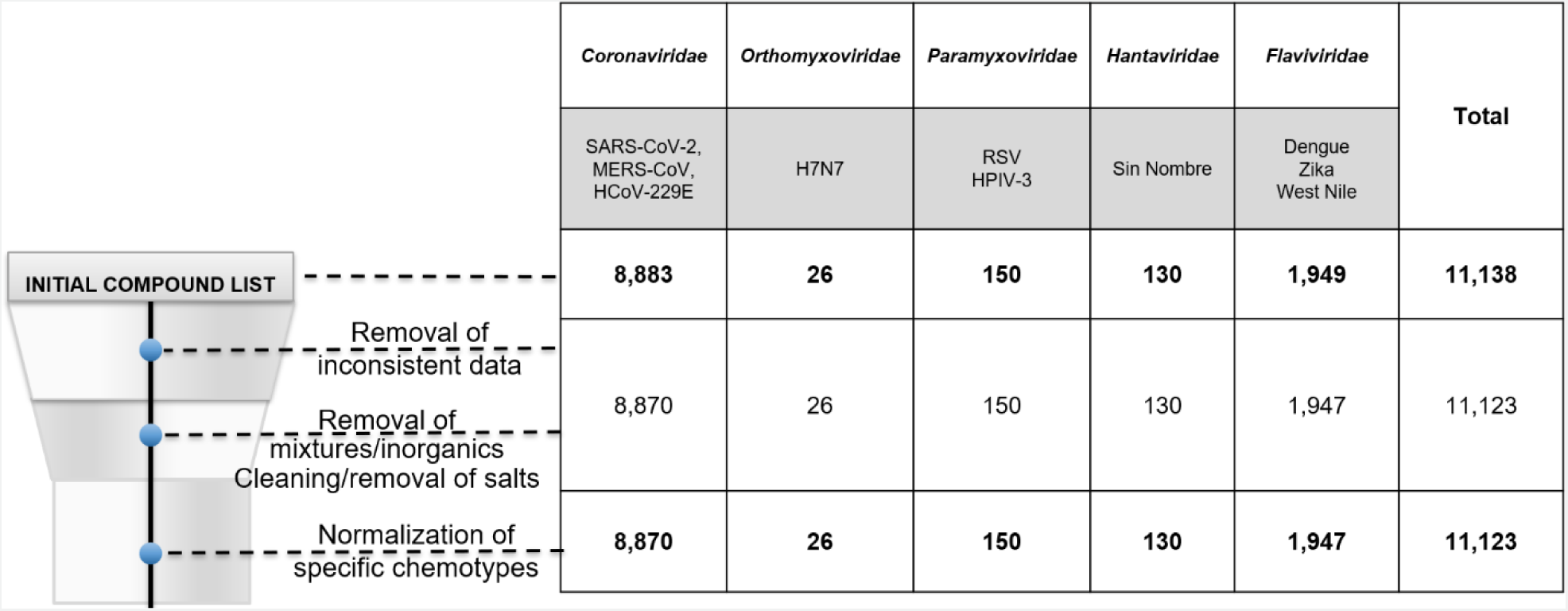
The effect of target-based assay data curation on reducing the resulting dataset sizes.

Note that at this step of data curation we intentionally kept duplicative compound records reflecting our objective to check whether the same compound showed similar activity against different viruses (i.e., had a potential to be a broad-spectrum agent). However, such chemically duplicative entries have been annotated in SMACC to facilitate their removal prior to the development of assay-specific QSAR models by the users of SMACC.

### Curated Phenotypic Data

Curated phenotypic testing entries in our database included assay data for 13 viruses in 5 viral families: *Coronaviridae* (SARS-CoV-2, MERS-CoV, HCoV-229E), *Orthomyxoviridae* (H1N2, H7N7), *Paramyxoviridae* (RSV, HPIV-3), *Phenuiviridae* (Sandfly Fever), and *Flaviviridae* (Dengue, Zika, Yellow Fever, Powassan, West Nile).

### Distribution of compound activity

The heatmap presented in **Figure 3** depicts the activity spectrum of all compounds tested in phenotypic assays for the viruses in our database. It is evident that many compounds were either inactive (80.6%) or untested. In contrast, the number of actives constituted 15.6% of the total number of entries, and the fraction of “true actives” with no conflicting assay results was just 6.48% (1,387 compounds). Unsurprisingly, the virus with the largest number of tested compounds was SARS-CoV-2 due to the many studies caused by the current pandemic, encompassing 61.6% of our phenotypic assay entries. Despite the enormous testing efforts, 94.76% of compounds were reported as inactive. Each virus had more inactive compounds in the dataset except Dengue (**Figure 3**).

**Figure 3.**
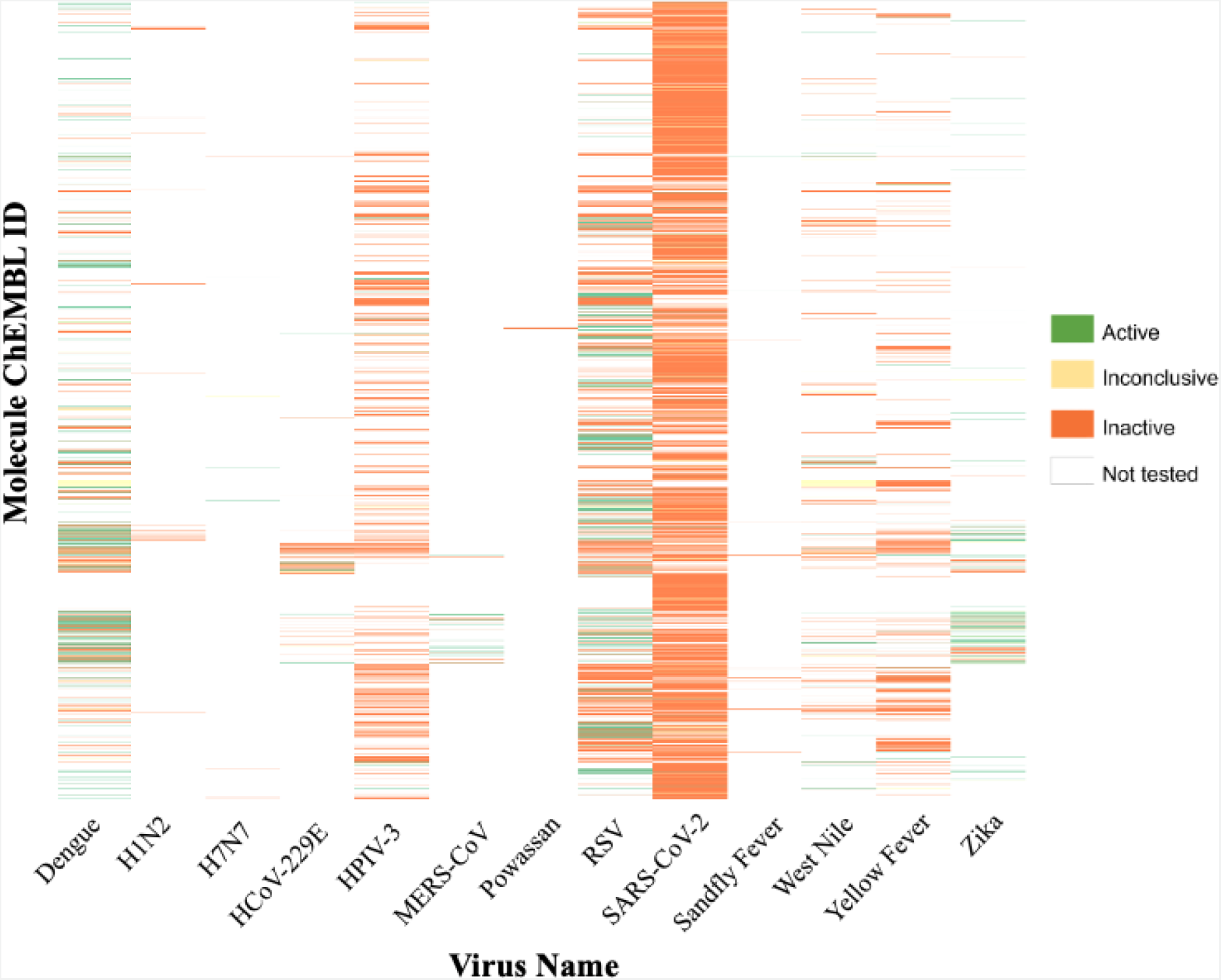
Activity heat map for 21,392 compounds tested in phenotypic assays for the 13 viruses selected for the SMACC database.

### Analysis of Cell Types

The 21,392 compounds integrated into our database were tested in 53 unique cell types. The most common cell types were Vero C1008 (37.6% of entries), Caco-2 (24.2%), Vero (9.1%), Huh-7 (4.6%), and Hep-2 (3.8%). The high propensity of testing in VeroC1008 cells is explained by a single assay screening against SARS-CoV-2 (8043 entries). Other cell types, such as Caco-2, were used for testing in multiple viruses and amongst various assays. Interestingly, Dengue virus had the greatest number of cell types tested (26), followed by RSV (17), and Zika virus (10) (**Figure 4**). Conversely, Vero cells were tested in the largest number of viruses (9) across 1,956 entries, followed by A549 (7 viruses), and Huh-7 cells (6 viruses).

**Figure 4.**
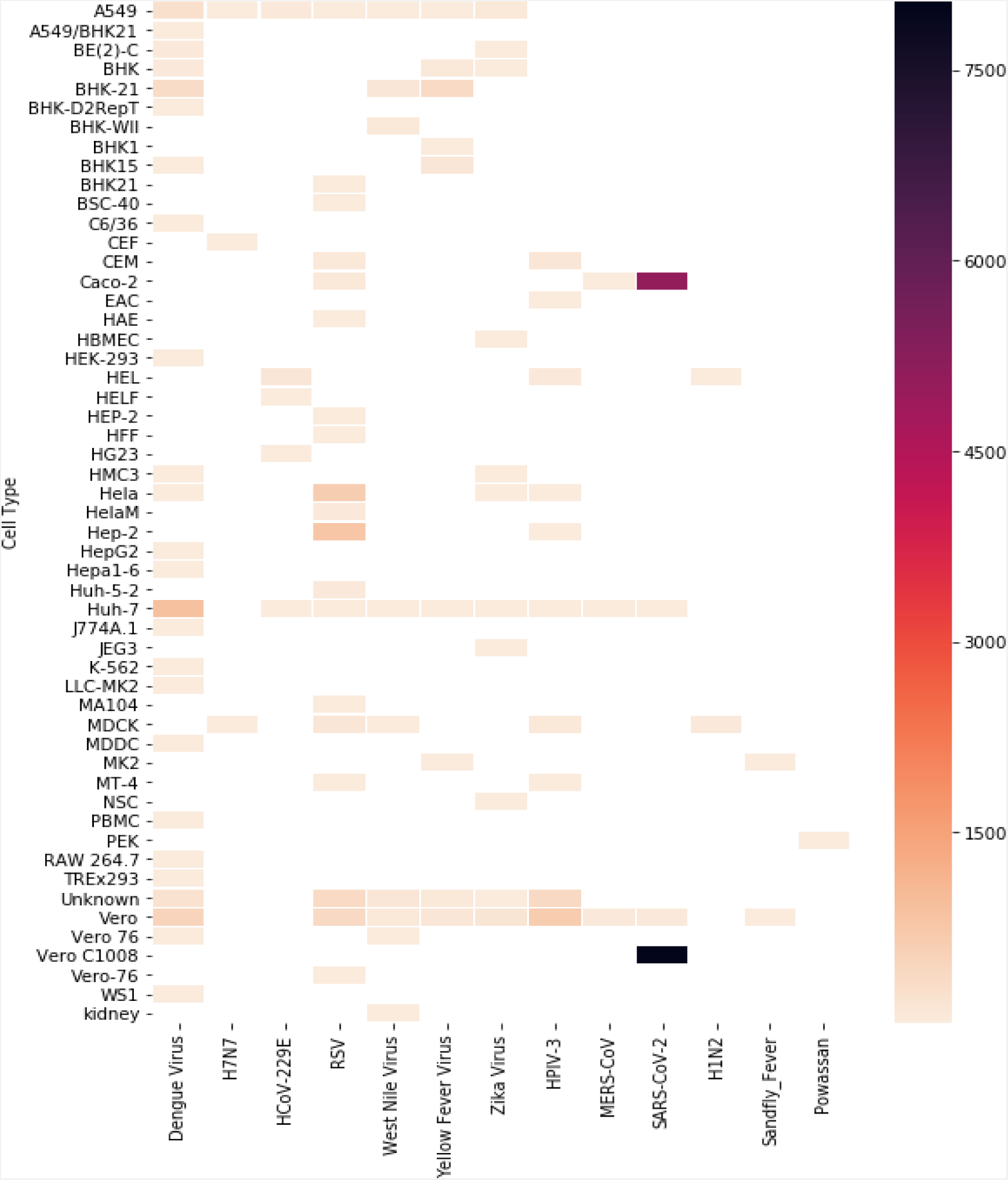
Heat map showing distribution of compounds tested in phenotypic screens for different cell types and different viruses.

### Stratifying compounds by assay type

We identified 27 compounds tested in the largest number of phenotypic assays in our database and further examined the effect of cell type on the resulting activity determination (Table S2). As expected, there were some inconsistencies in the activities determined when stratifying compounds by the virus they were tested against and the cell type for that virus and then comparing their activities. The data for these 27 compounds have been recorded in a matrix with 27 respective columns and 146 rows (Table S2). Unknown cell types and inconclusive activity results were ignored. We identified 26 assay results when a compound was tested in the same virus and cell line but showed conflicting results. In another 19 cases a compound was tested against the same virus but in different cell lines, and had different results. In contrast, for 10 cases, we observed completely consistent activity testing results (in 2+ entries) for a compound assayed in the same virus in the same cell line. We also observed 22 cases of consistent activity when compounds were tested for the same virus but in multiple cell lines. There were also many cases reporting a compound tested for a single virus and multiple cell types. In this case, we only analyzed whether or not the activities reported for each cell line were consistent. In summary, we observe that the choice of cell type can influence the outcome of the assay, an observation reported previously ^17^ and thus, the annotation of a compound as active or inactive against any virus should be always reported strictly in the context of the specific underlying assay. Consequently, integration of data across multiple cell line, for instance, to increase the size of the data for QSAR model development, should be done with care, i.e., only when the evidence exists that compounds show similar activities when tested in different cell lines.

### Identifying compounds active in multiple assays

Our efforts to extract and consolidate data on compounds tested in antiviral assays, as reported in ChEMBL, revealed that identifying truly active compounds is complex and requires careful curation of the available assay results. We first selected a subset of compounds from our database that were tested in assays against two or more viruses. As we described in Methods, to analyze the multi-viral activity we annotated each compound with one of three possible activity calls. A compound was considered active if it was recorded as active when tested in all assays, inactive if it was inactive in all assays, and considered inconclusive if it was active in some assays and inactive in others (Table 1), if the assay result was reported with an ambiguous operator (>or<), or if the compounds activity was not successfully determined by the assay. From this, we created an intermediate table where each compound occupied one row, and the columns contained concatenated lists of every virus the compound was reported active or inactive against. We systematically analyzed this matrix to identify compounds that we considered to be true actives, i.e., the compound was reported active in all assays in which it was tested.

We report the eight most promising compounds resulting from this analysis in **Table 3**. Here our top compound is CHEMBL4437334 with activity against Dengue, West Nile, Yellow Fever, and Zika. It is a research compound not yet progressed into any clinical trial, which is true for many compounds of this list including CHEMBL4454780 (active against RSV, MERS-CoV, Dengue, and Zika), CHEMBL2016757 (active against RSV, HPIV-3, and Dengue), CHEMBL4544911 and CHEMBL4562509 (active against Dengue, West Nile, and Yellow Fever). Three named compounds were identified from our search: 6-azauridine (active against RSV, West Nile, Dengue); amodiaquine (active against SARS-CoV-2, Dengue, Zika), which is an approved drug for malaria; and brequinar (active against Dengue, West Nile, and Yellow Fever), which is currently in Phase I clinical trials for treatment of acute myeloid leukemia.^18–20^

**Table 3.**
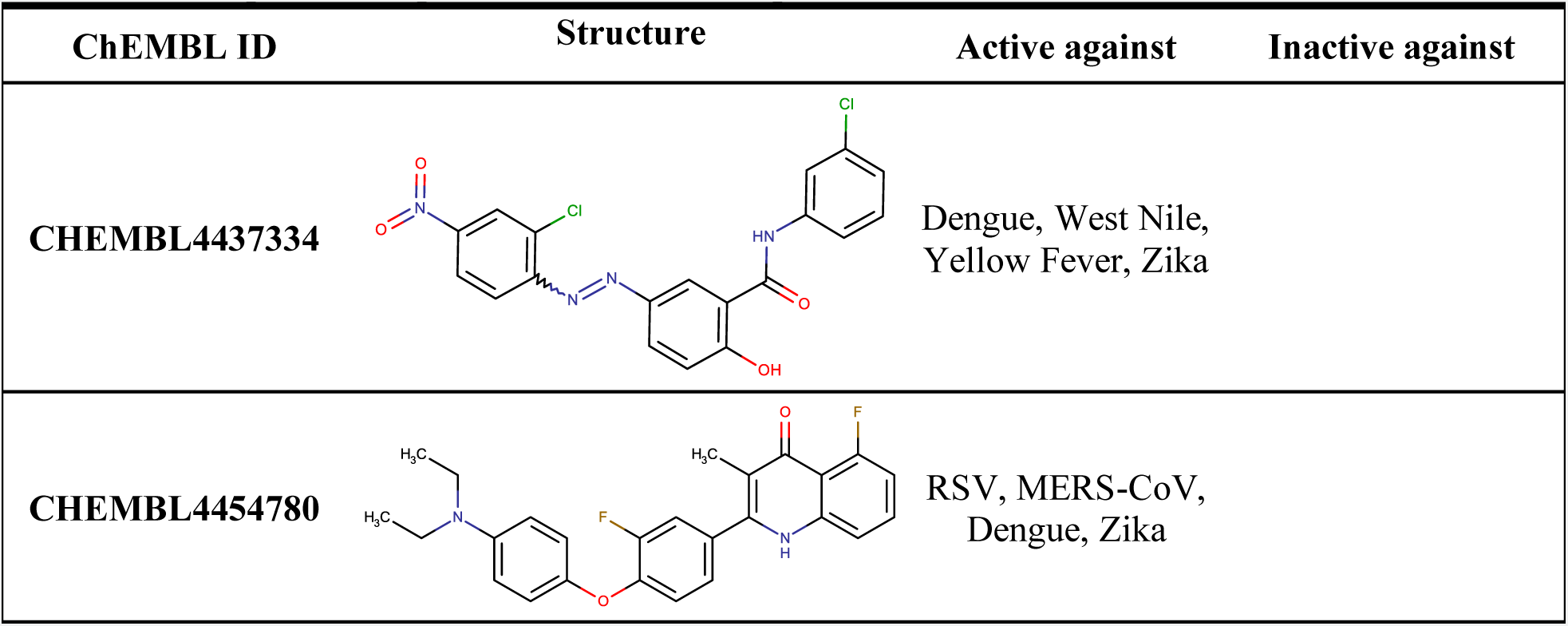
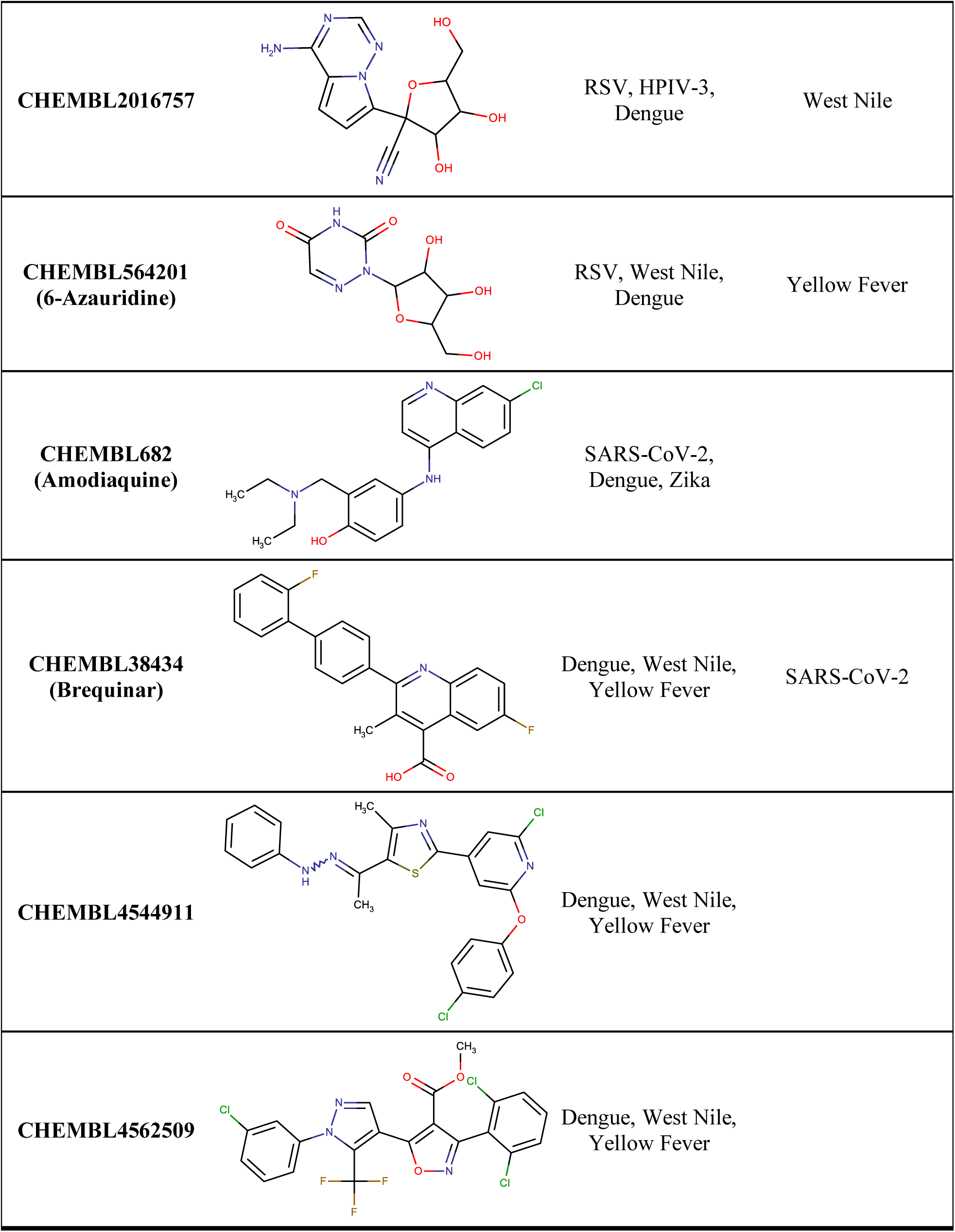
Example of compounds active in multiple viruses.

| ChEMBL ID | Structure | Active against | Inactive against |
| --- | --- | --- | --- |
| CHEMBL4437334 |  | Dengue, West Nile, Yellow Fever, Zika |  |
| CHEMBL4454780 |  | RSV, MERS-CoV, Dengue, Zika |  |
| **CHEMBL2016757** |  | RSV, HPIV-3,<br>Dengue | West Nile |
| **CHEMBL564201**<br>(6-Azauridine) |  | RSV, West Nile,<br>Dengue | Yellow Fever |
| **CHEMBL682**<br>(Amodiaquine) |  | SARS-CoV-2,<br>Dengue, Zika |  |
| **CHEMBL38434**<br>(Brequinar) |  | Dengue, West Nile,<br>Yellow Fever | SARS-CoV-2 |
| **CHEMBL4544911** |  | Dengue, West Nile,<br>Yellow Fever |  |
| **CHEMBL4562509** |  | Dengue, West Nile,<br>Yellow Fever |  |

Interestingly, brequinar also recently underwent phase II clinical trials against SARS-CoV-2.^21^ While the clinical trial was not successful using brequinar alone, research on brequinar drug combinations have found this drug highly effective in combination with remdesivir or molnupiravir.^22^ Research suggests that brequinar’s antiviral activity against SARS-CoV-2 is through inhibition of the host cell dihydroorotate dehydrogenase (DHODH) rather than being a direct acting antiviral. The combination of a nucleobase antiviral and DHODH, or other compound that impacts de novo nucleotide synthesis would in effect increase the nucleobase antiviral cellular concentration thereby increasing the rate of incorporation into the viral synthesized RNA, in theory. This approach has been shown to be effective against multiple viruses *in vitro*, for example, brequinar has been shown to inhibit dengue, enterovirus, and Ebola viruses through this same, host-targeted mechanism.^23–25^ Given the reported activity of brequinar against three *flaviviruses* (Dengue, West Nile, Yellow Fever) in phenotypic assays, we hypothesize the assays were detecting the human dihydroorotate dehydrogenase inhibition, rather than inhibition of a viral target. As such, a future release of SMACC will include an analysis of all phenotypic assays, host-target assays, and the overlap between the phenotypic assays and the host-target assays. It is our hope this analysis will help develop hypotheses for, and identify, potential host-targeting broad-spectrum antiviral drugs.

There were 55 more compounds active against at least two viruses (**Table S3**). Of these, seven compounds were active against different viral families: three were active against *Paramyxoviridae* (RSV, HPIV-3); two were active against *Coronaviridae* (MERS-CoV, HCoV-229E); and 43 against any two of our *Flaviviridae* viruses (Dengue, Zika, Yellow Fever, Powassan, and West Nile). We also identified 1,324 compounds active against one virus.

### Cluster analysis of active compounds

The structural clustering of all compounds tested in phenotypic cell-based assays revealed ten clusters (**Figure 6**). The top BSA compounds, CHEMBL4437334 and CHEMBL4454780, active against four different viruses, are in clusters #7 and #5, respectively. The subcluster containing CHEMBL4437334 (cluster #7) has 847 compounds. Among them, some nearest neighbors of CHEMBL4437334 (**Figure 7**) were active against SARS-CoV-2 and could be further tested against a panel of flaviviruses (Dengue, West Nile, Yellow Fever, and Zika). The subcluster of CHEMBL4454780 (cluster #5) contains 1,406 compounds; nearest neighbors of CHEMBL4454780 are presented in **Figure 7**. Compounds CHEMBL1197690, CHEMBL3581155, and CHEMBL7568 were active against one or two flaviviruses and could be further tested against additional flaviviruses and viruses from other families like RSV and MERS-CoV. Likewise, CHEMBL4303559 was only tested and active against SARS-CoV-2 and could be tested against members of *Flaviviridae* and other coronaviruses such as MERS-CoV.

**Figure 6.**
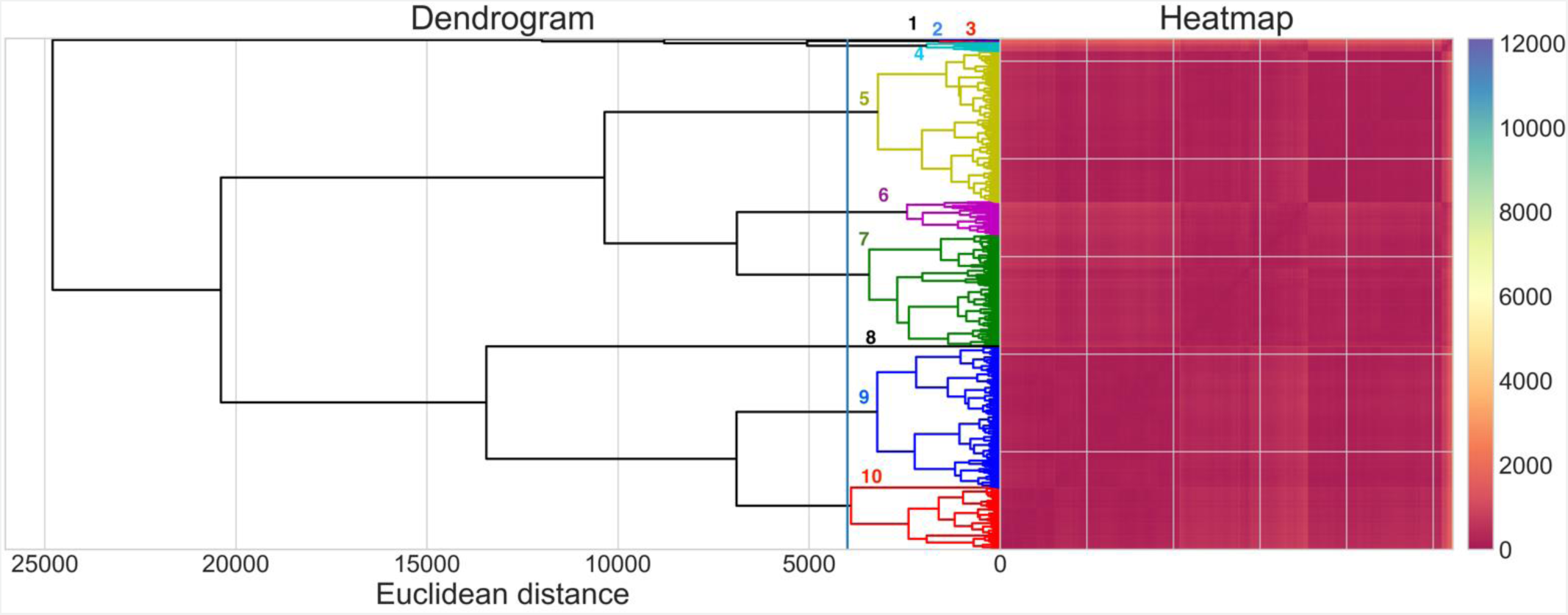
Clustering of compounds from the phenotypic cell-based assays by chemical structure. The colors from the heatmap are based on the Euclidean distances between compounds. Colors nearer to dark red indicate a shorter distance between molecules.

**Figure 7.**
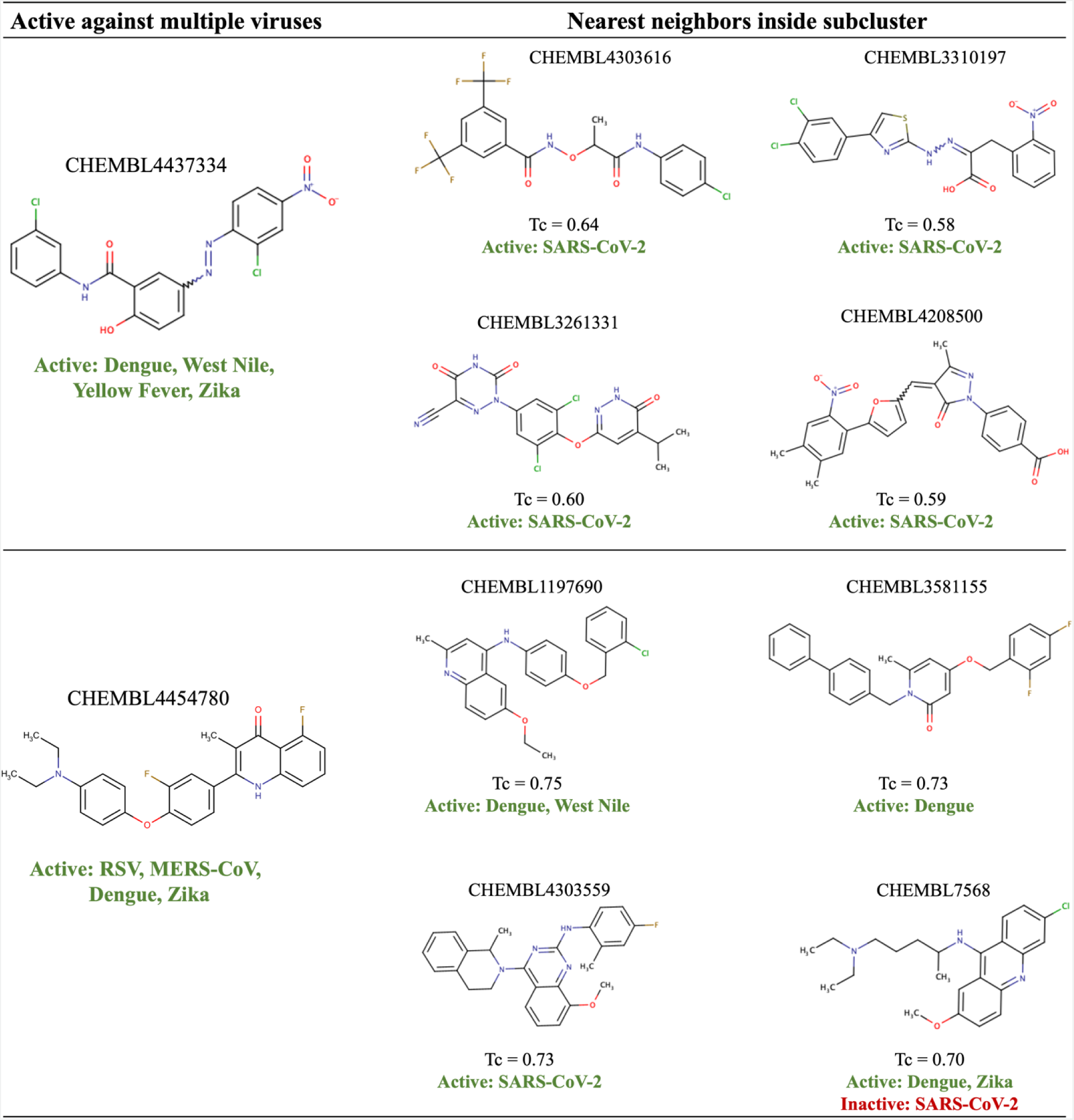
Examples of compounds similar to experimental broad-spectrum hits that could be further tested against multiple viruses of interest.

### Target-Based Data

Curated target-based testing entries (11,123) in our database include assay data for ten viruses in five viral families: *Coronaviridae* (SARS-CoV-2, MERS-CoV, HCoV-229E), *Orthomyxoviridae* (H7N7), *Paramyxoviridae* (RSV, HPIV-3), *Phenuiviridae* (Sin Nombre), and *Flaviviridae* (Dengue, Zika, West Nile).

### Activity of Compounds

Using the activity calls based on the assay results as defined in Methods, most compounds (89.8%) were inactive (**Figure 8**). Many compounds inactive against SARS-CoV-2 were tested because of the recent multiple testing campaigns including drug repurposing screenings due to current pandemic. While SARS-CoV-2 was the most tested virus, three flaviviruses were also well represented in the database (Dengue, West Nile, and Zika); however, as the number of compounds tested increased there was a decrease in the fraction of the active compounds for these viruses. Overall, active compounds represented only ∼9.9% of our total dataset where 5.78% (644 compounds) were “true actives”.

**Figure 8.**
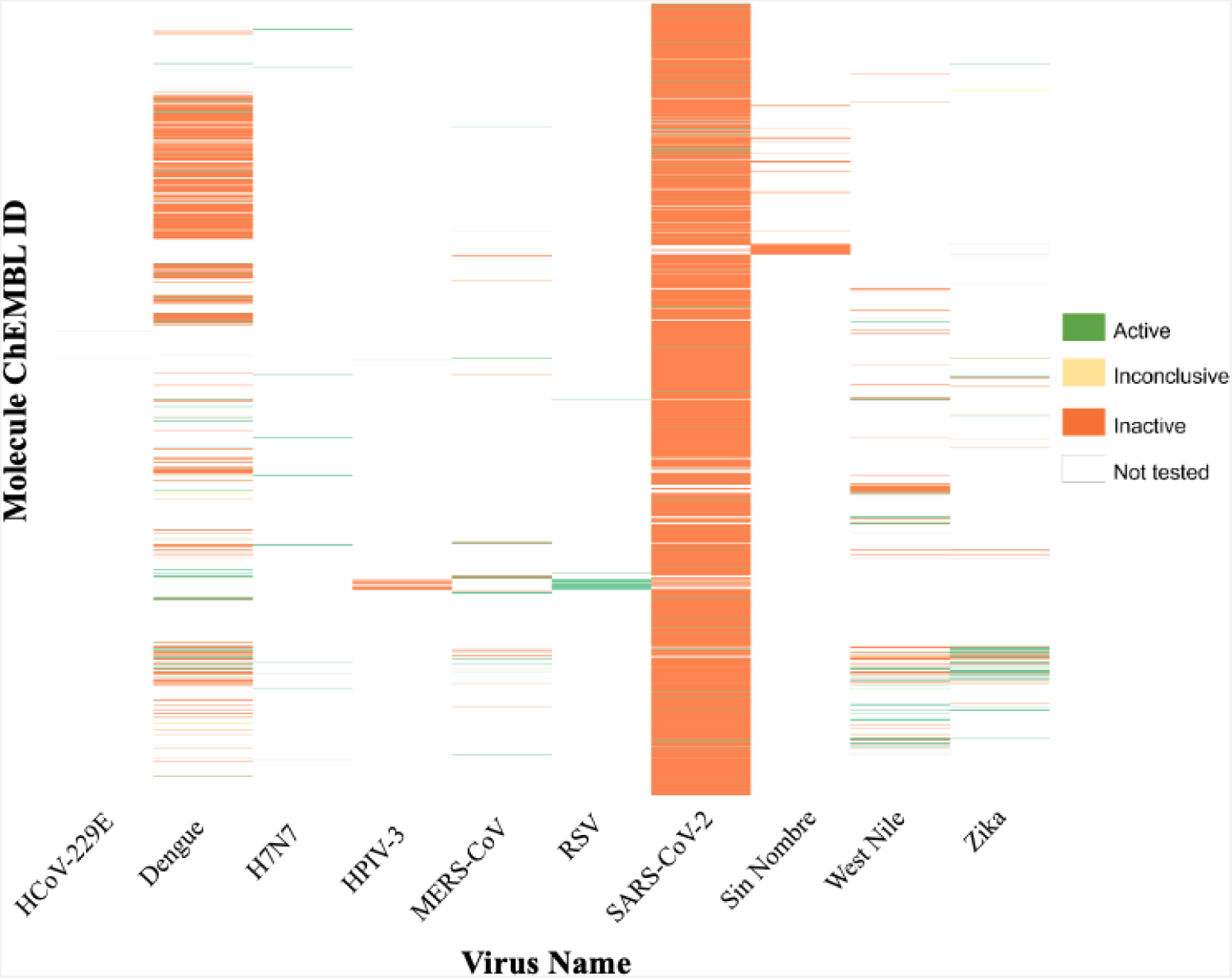
Activity heat map for 11,123 compounds tested in target-based assays for the 10 viruses selected for the SMACC database.

### Analysis of Targets

The Main Protease (3CLpro) of *Coronaviridae* was, unsurprisingly, the most studied target (78.8% of the entries), followed by NS2B-NS3 Protease of *Flaviviridae* (16.2%), NS5 of *Flaviviridae* (1.26%), Integrin alpha-V/beta-3 of *Hantaviridae* (1.17%), and Fusion glycoprotein F0 of *Paramixoviridae* (1.1%). Interestingly, the virus with the greatest number of targets tested (five) was MERS-CoV and was tested against the spike protein, RDRP, Nucleocapsid protein, M^pro^, and PL^pro^.

### Analysis of Compounds

We followed the same approach for analyzing the target-based dataset for BSA activity, as was taken for the phenotypic dataset. In this case, the intermediate table included a row for each compound, and the columns were concatenated lists of every virus and target the compound was reported active or inactive against. Our analysis identified 16 compounds active against two viruses at the protein target level (**Table 4**). Two of these compounds (CHEMBL4544781 and CHEMBL4522602) were active against targets from two different viral families (Zika’s NS5 and MERS-CoV’s RDRP), whereas the others were active against two flaviviruses NS2B-NS3 Protease. We also identified 628 compounds active against one virus.

**Table 4.**
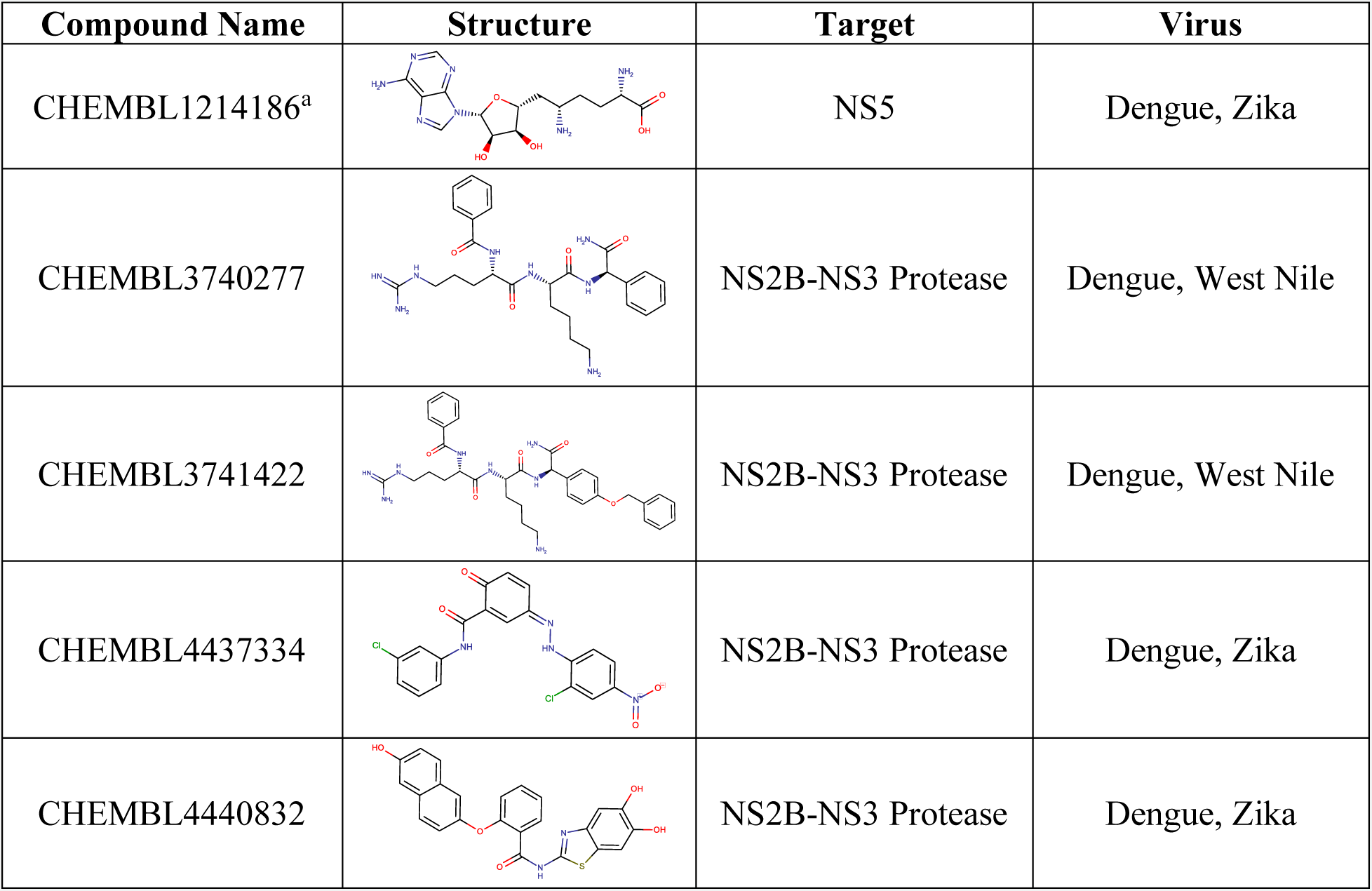
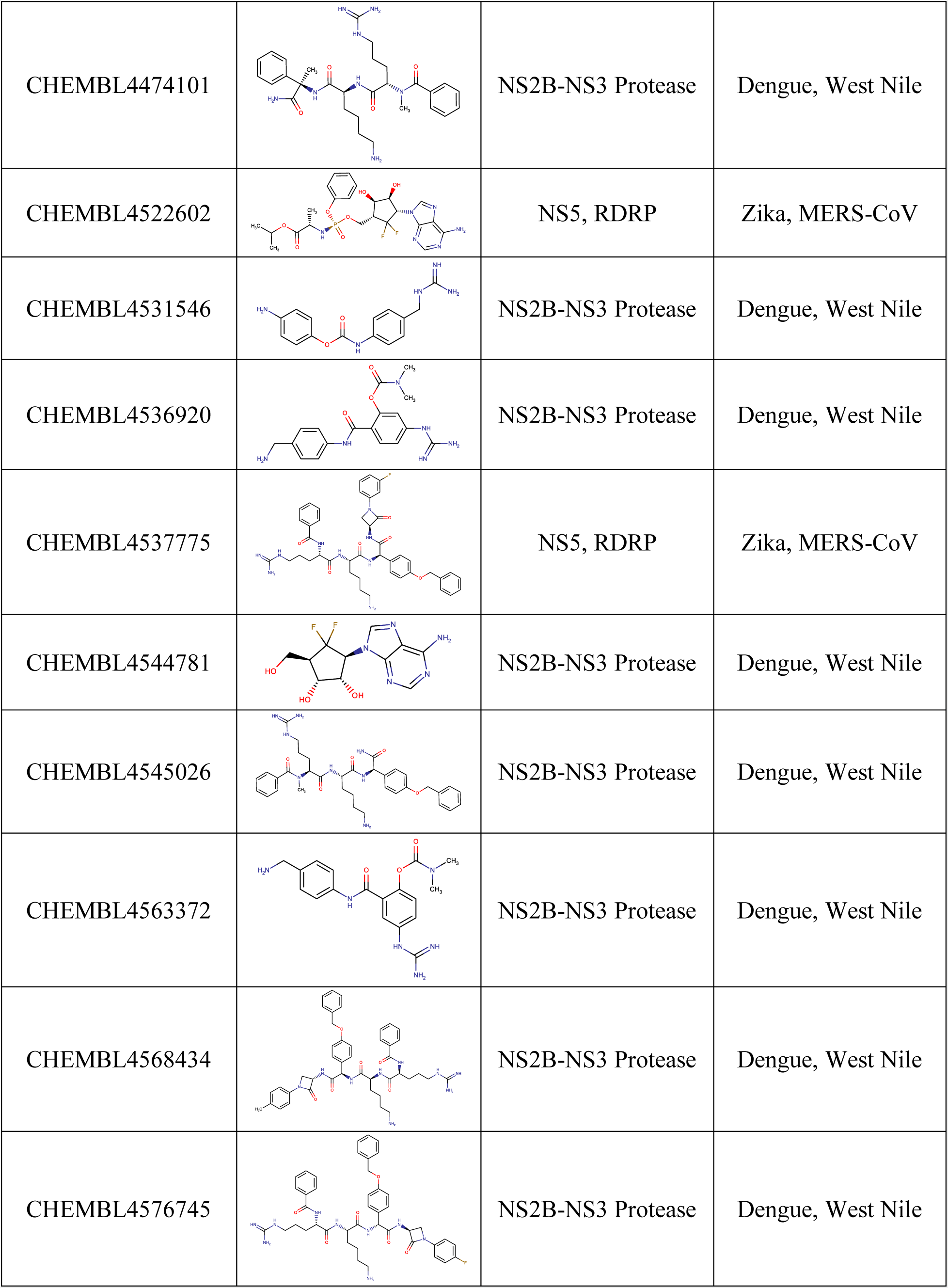
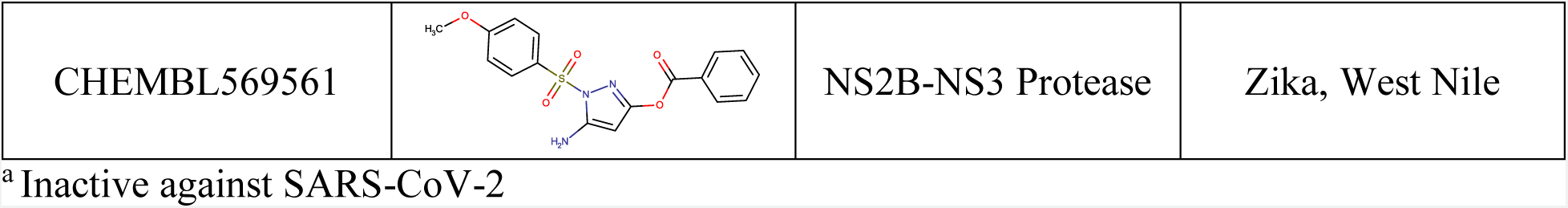
Compounds active against different viruses in target-based assays.

| Compound Name | Structure | Target | Virus |
| --- | --- | --- | --- |
| CHEMBL1214186<sup>a</sup> |  | NS5 | Dengue, Zika |
| CHEMBL3740277 |  | NS2B-NS3 Protease | Dengue, West Nile |
| CHEMBL3741422 |  | NS2B-NS3 Protease | Dengue, West Nile |
| CHEMBL4437334 |  | NS2B-NS3 Protease | Dengue, Zika |
| CHEMBL4440832 |  | NS2B-NS3 Protease | Dengue, Zika |
| CHEMBL4474101 |  | NS2B-NS3 Protease | Dengue, West Nile |
| CHEMBL4522602 |  | NS5, RDRP | Zika, MERS-CoV |
| CHEMBL4531546 |  | NS2B-NS3 Protease | Dengue, West Nile |
| CHEMBL4536920 |  | NS2B-NS3 Protease | Dengue, West Nile |
| CHEMBL4537775 |  | NS5, RDRP | Zika, MERS-CoV |
| CHEMBL4544781 |  | NS2B-NS3 Protease | Dengue, West Nile |
| CHEMBL4545026 |  | NS2B-NS3 Protease | Dengue, West Nile |
| CHEMBL4563372 |  | NS2B-NS3 Protease | Dengue, West Nile |
| CHEMBL4568434 |  | NS2B-NS3 Protease | Dengue, West Nile |
| CHEMBL4576745 |  | NS2B-NS3 Protease | Dengue, West Nile |
| CHEMBL569561 |  | NS2B-NS3 Protease | Zika, West Nile |

<sup>a</sup> Inactive against SARS-CoV-2

Structural clustering of all compounds revealed 11 clusters (**Figure 9**). CHEMBL4544781 was active against targets from different viral families (NS5 of Zika Virus and RDRP of MERS-CoV) and is in cluster #7 along with 867 other compounds; nearest neighbors of CHEMBL4544781 are presented in **Figure 10**. CHEMBL1630221 was only tested (and active) against the NS5 Polymerase of Zika Virus and could be further tested against other polymerases from other flaviviruses and the RNA-Dependent RNA Polymerase (RdRP) of MERS-CoV. Three other nearest neighbors of CHEMBL4544781 were only tested (and active) against SARS-CoV-2 Main Protease (M^pro^). These compounds could be further tested against polymerases of Zika and MERS-CoV.

**Figure 9.**
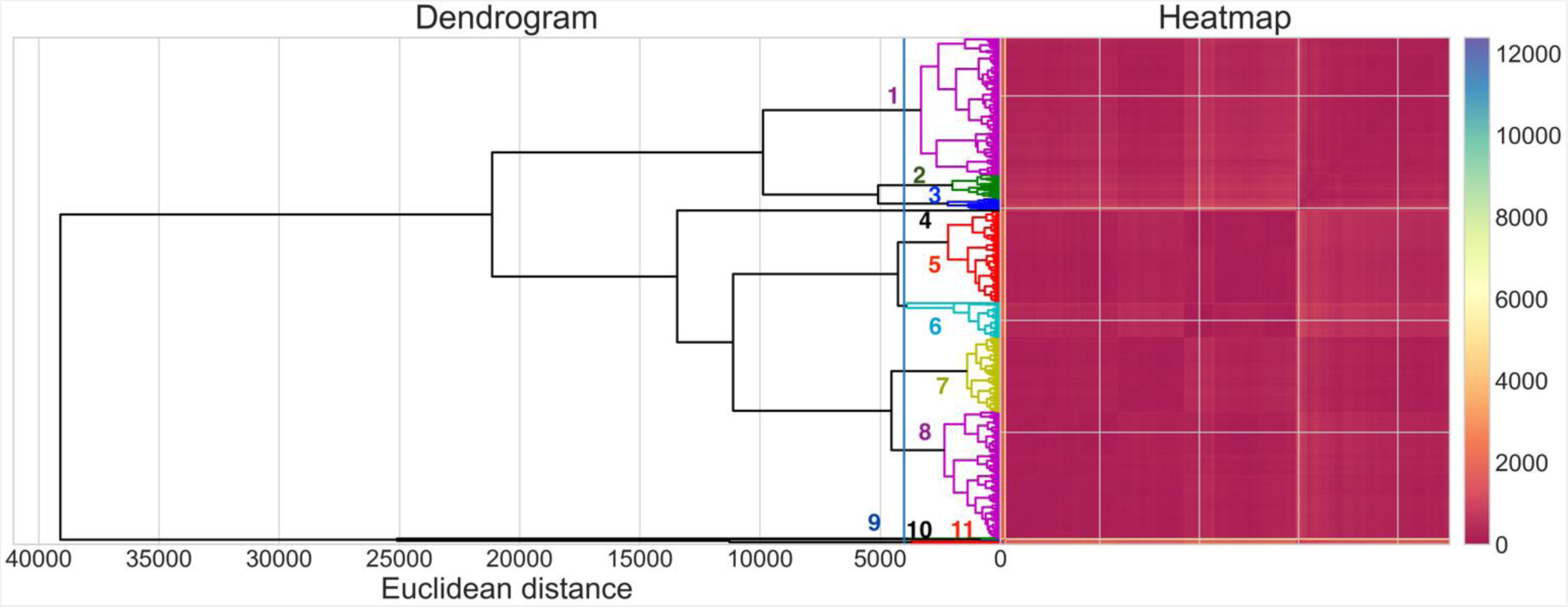
Clustering analysis of compounds from the target-based assays. The colors from the heatmap are based on the Euclidean distances between compounds. Colors nearer to dark red indicate a shorter distance between molecules.

**Figure 10.**
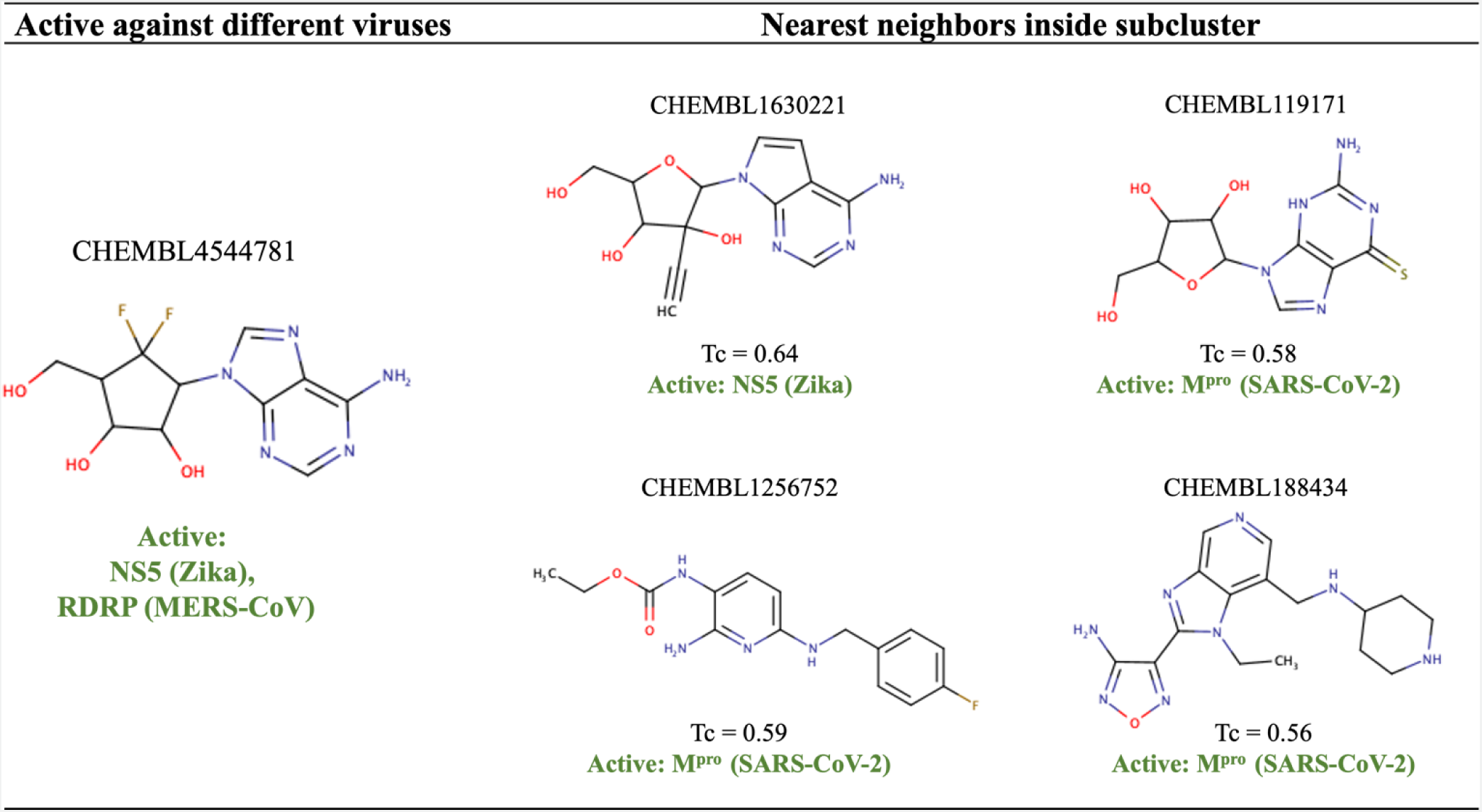
Examples of compounds that could be further tested against different viral targets of interest due to their chemical similarity to an active molecule with multiple antiviral activity.

### Concordance between the phenotypic and target-based data

We analyzed the concordance between the 5,934 compounds tested in both phenotypic and target-based assays to: (i) expand our list of hits by identifying potentially promising compounds that may not have been tested yet, and (ii) hypothesize the mechanism of actions of compounds active in a virus in a live cell and a complementary viral target. Our systematic analysis of the assay results indicated that 35 compounds were active in at least one phenotypic and one target-based assay (**Table S4**, Supplementary Material). In many cases, the active calls were within the same viral family. For example, CHEMBL4522006 was active against Dengue Virus in a phenotypic assay, and active against the Dengue NS2B-NS3 Protease in a target-based assay. Our data strongly supports the hypothesis that CHEMBL4522006 is active against Dengue virus in the live cell assay by inhibiting its NS2B-NS3 Protease, which supports the use of the protease assays for future experimental and computational structure-activity relationship studies. Promising potential BSA compounds, including CHEMBL4522006, are summarized in **Table 5**.

**Table 5.**
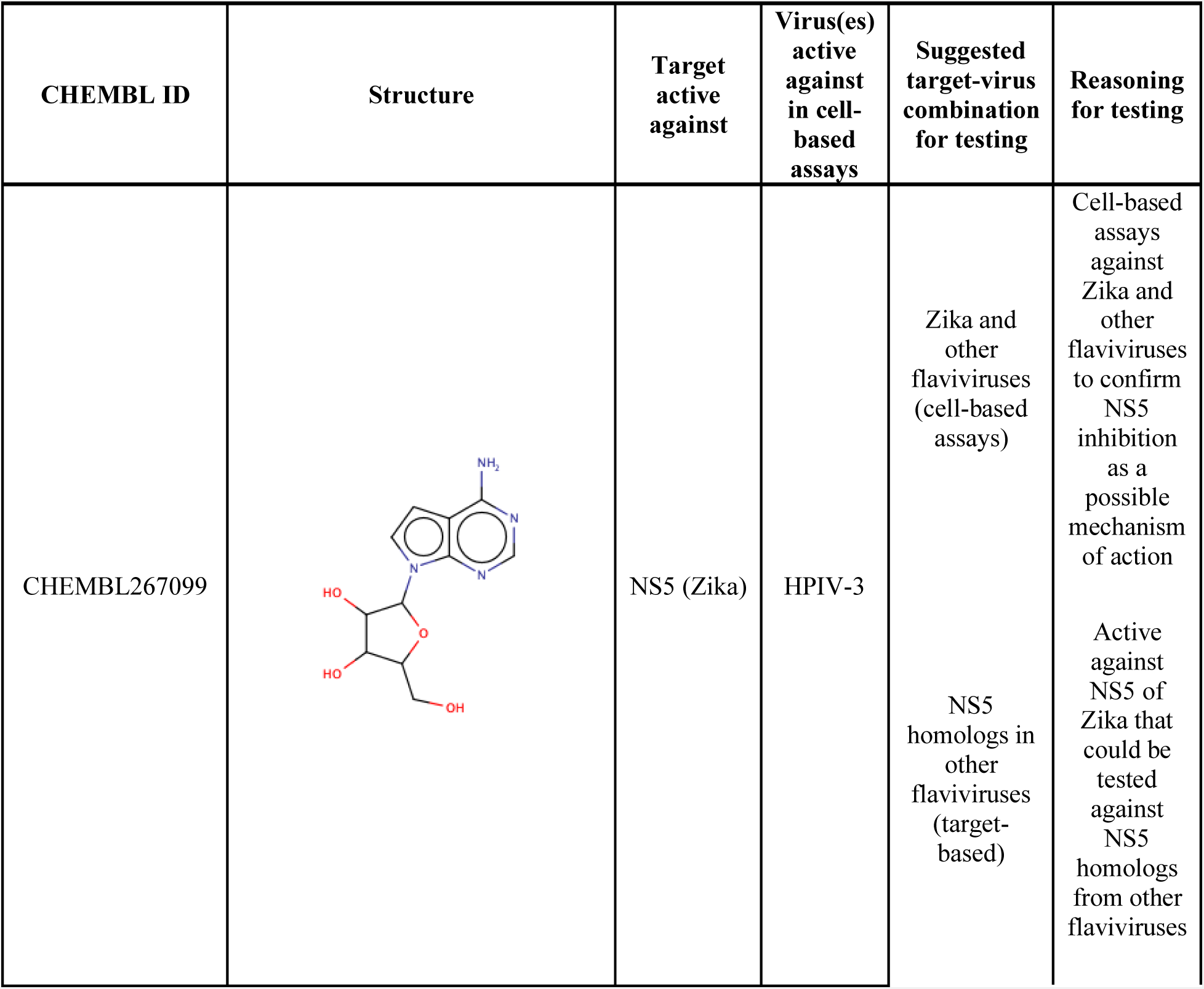
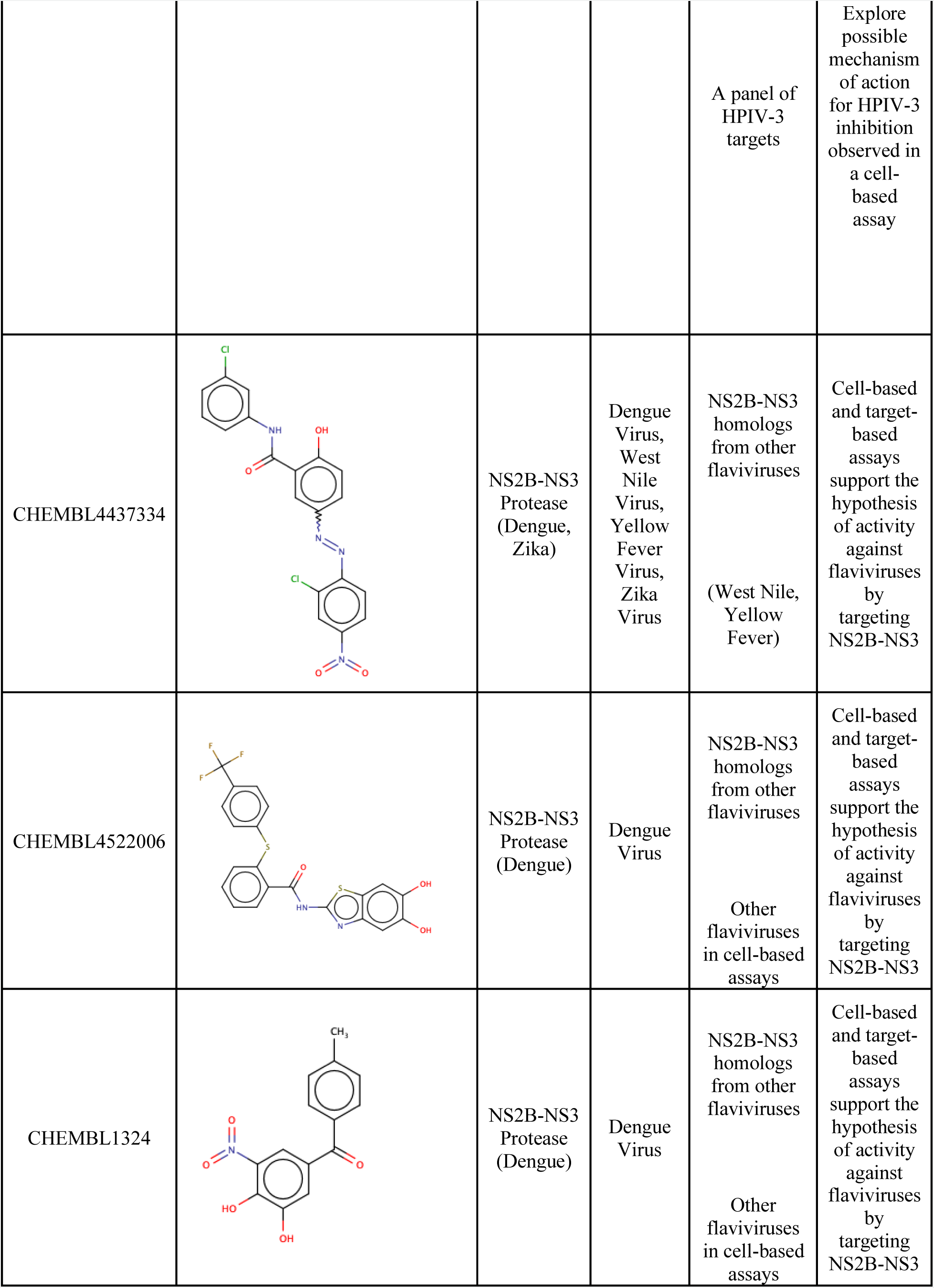
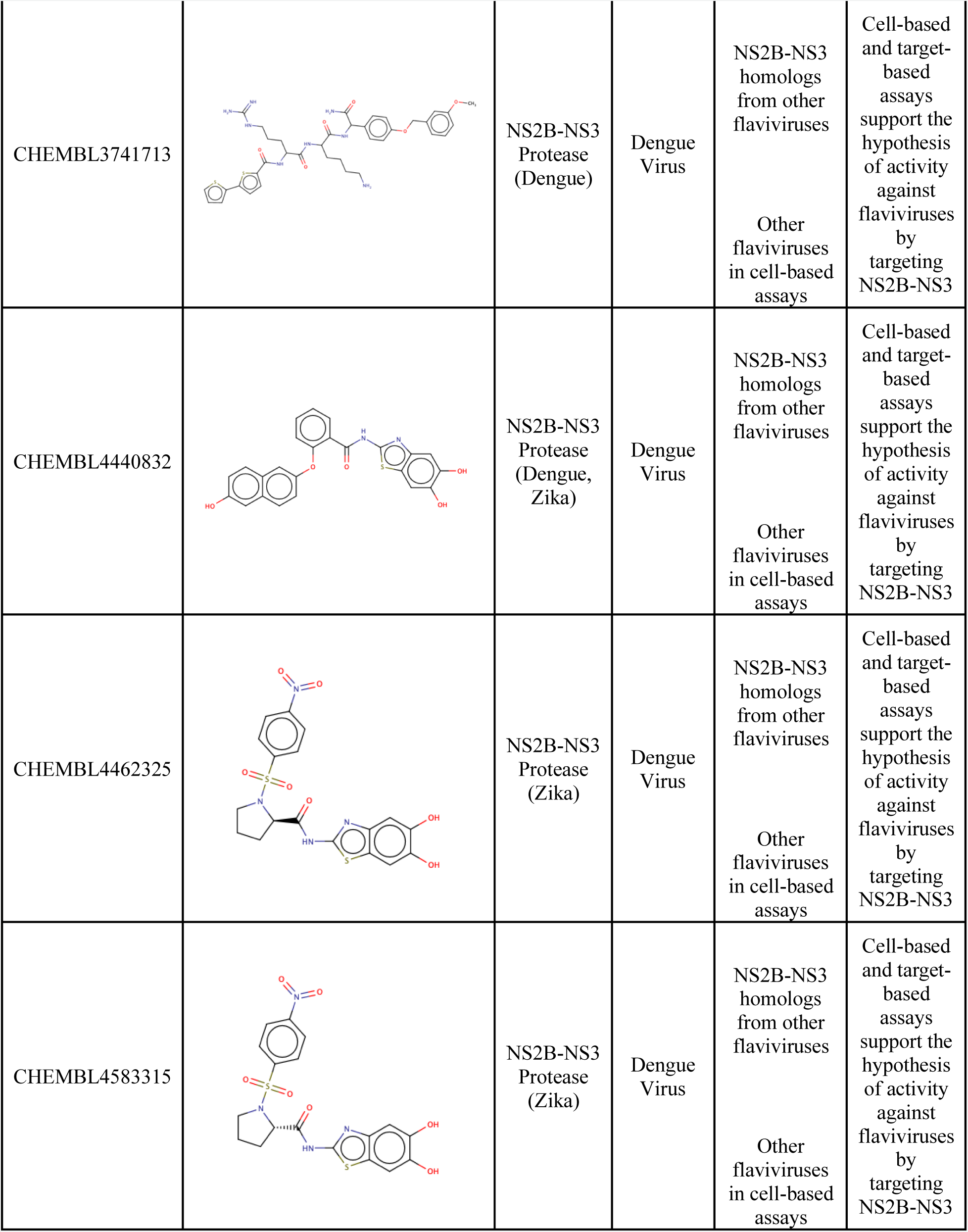
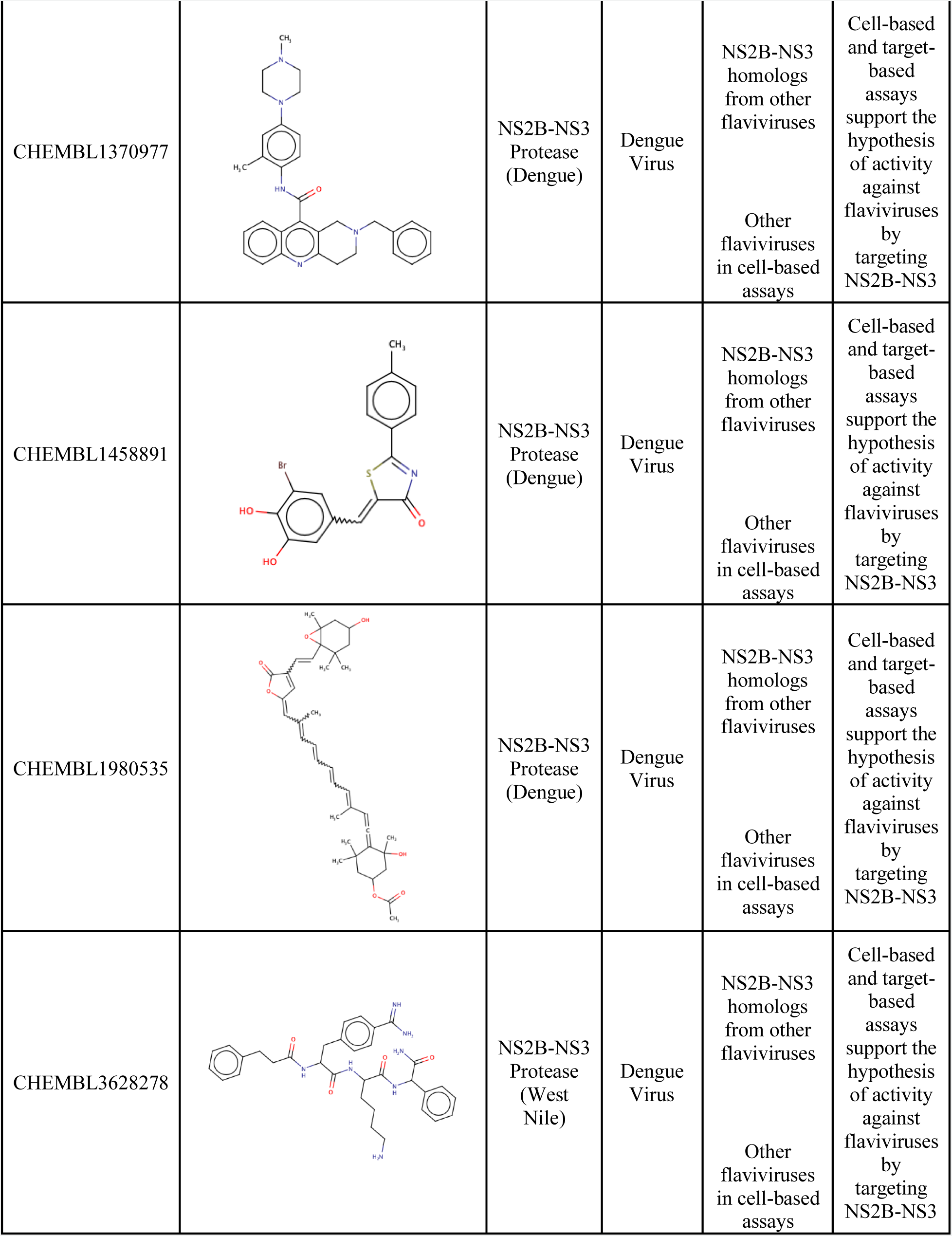
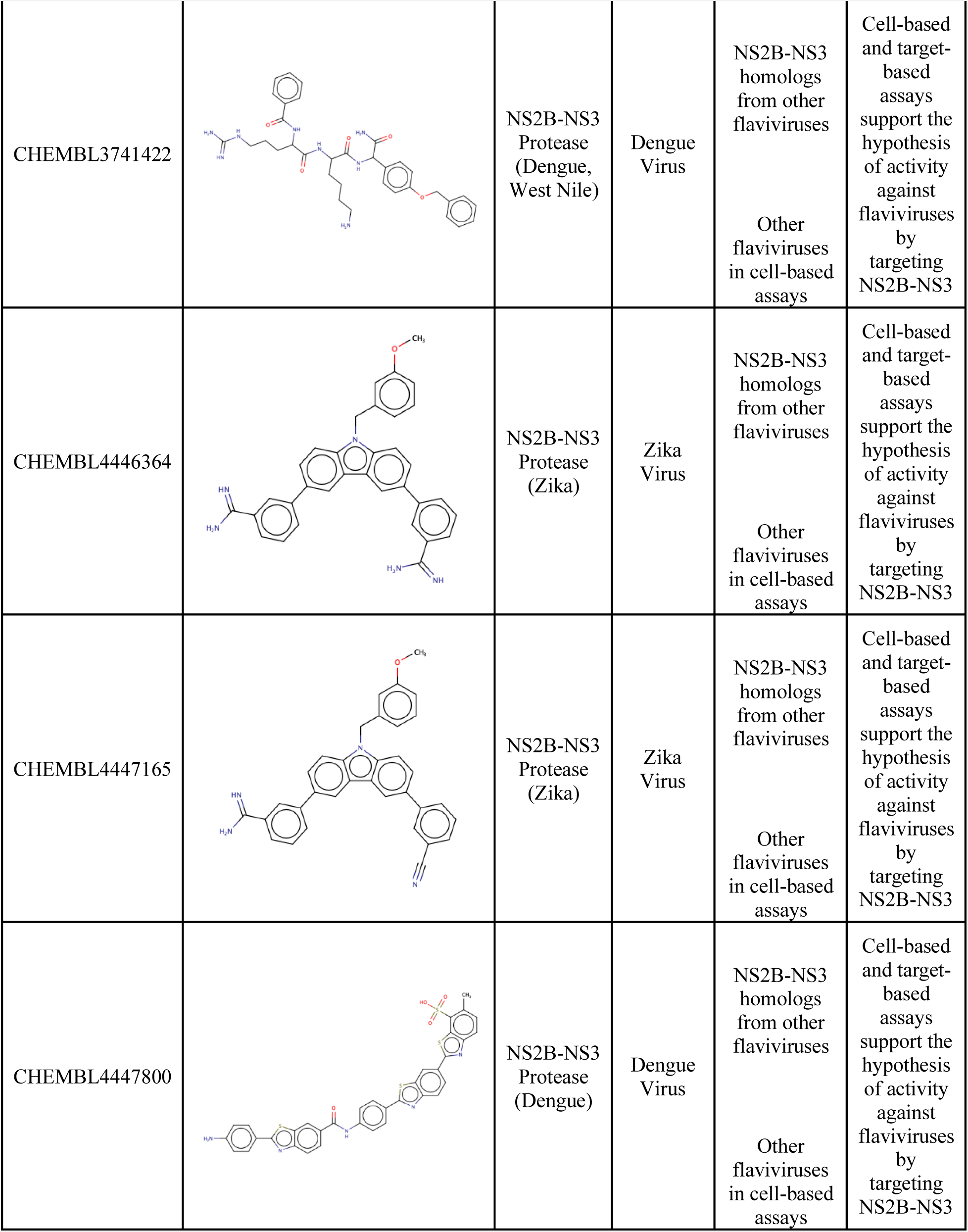
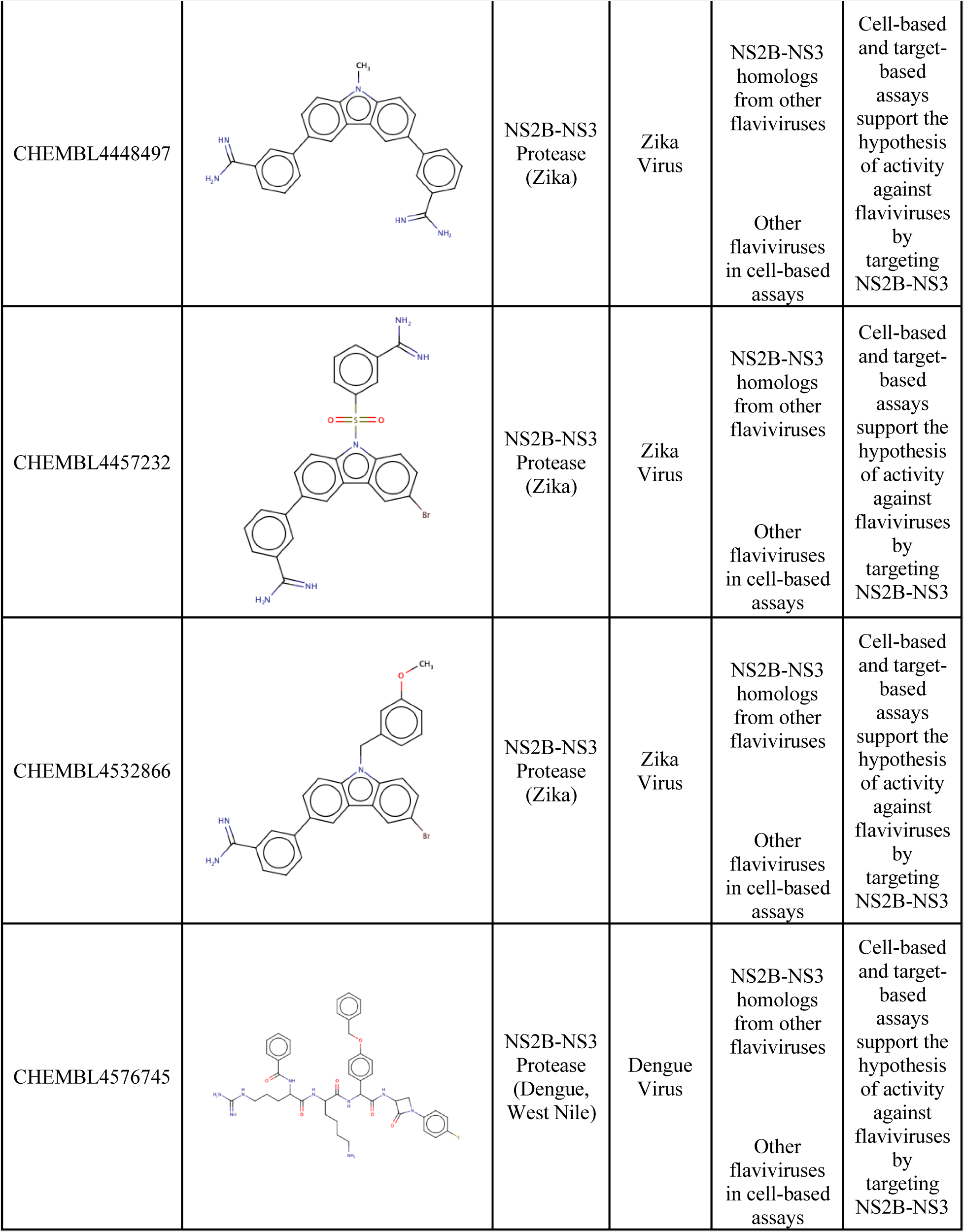
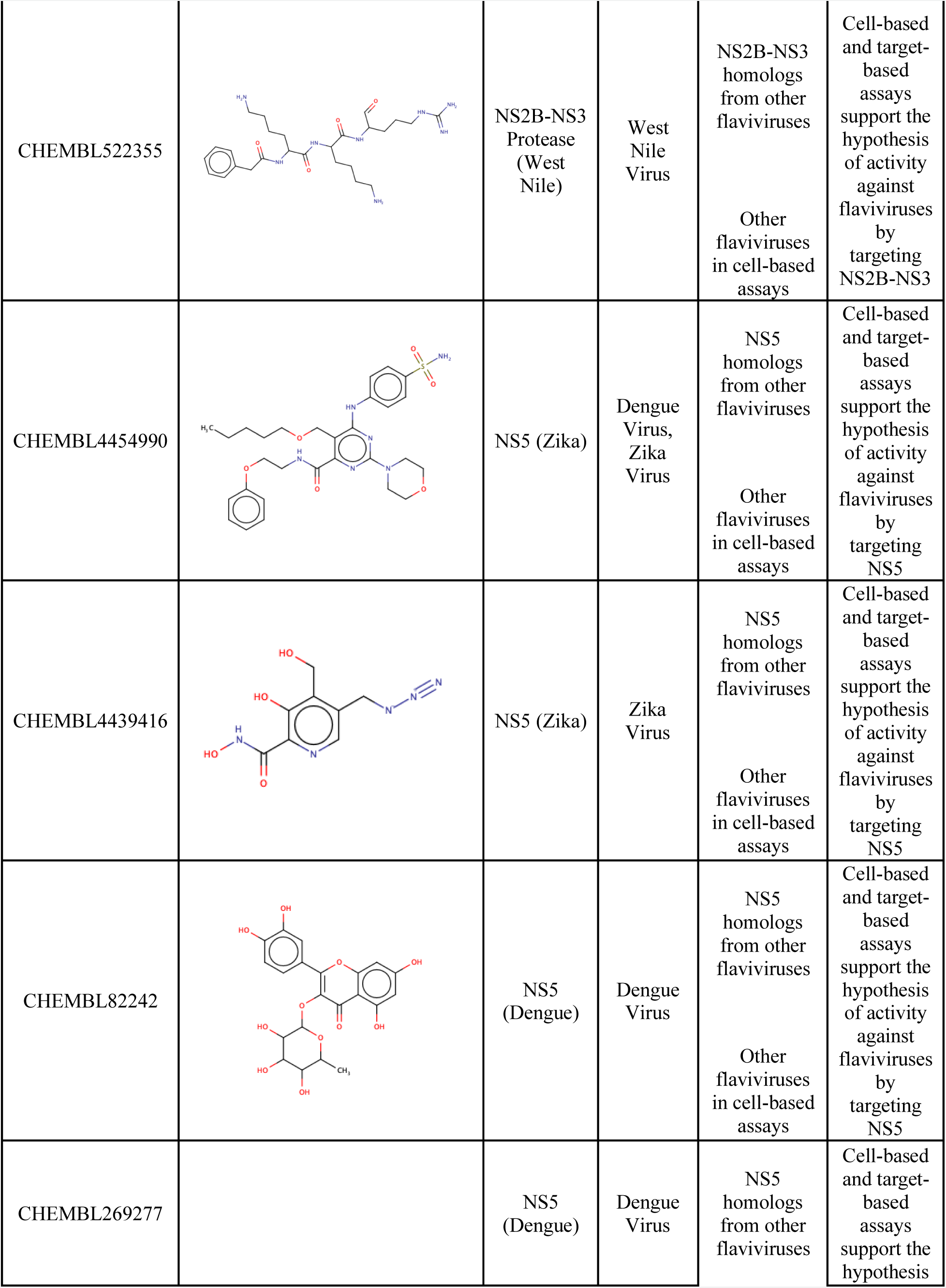
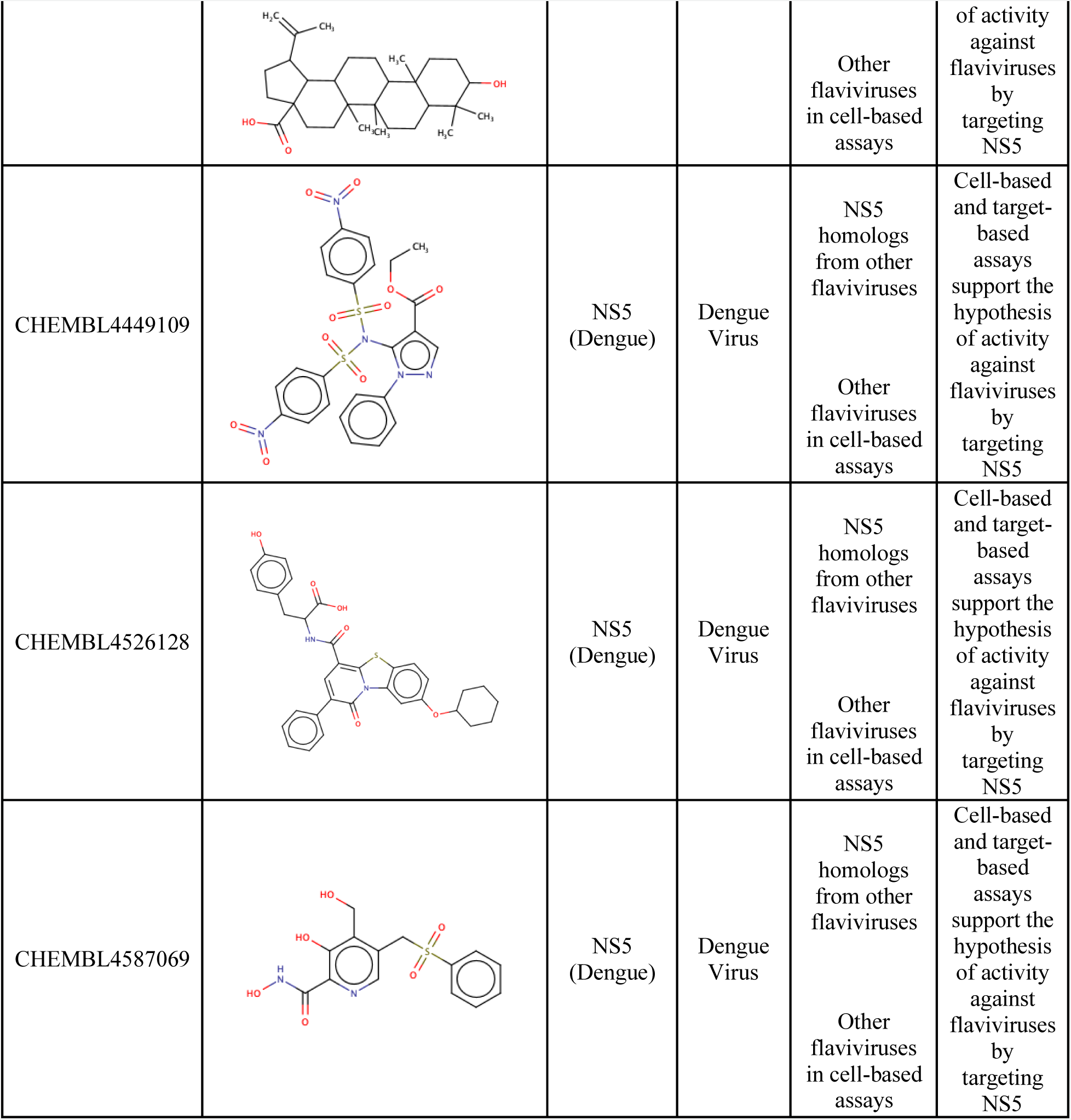
Selection of compounds nominated for experimental testing as potential BSA agents

| CHEMBL ID | Structure | Target active against | Virus(es) active against in cell-based assays | Suggested target-virus combination for testing | Reasoning for testing |
| --- | --- | --- | --- | --- | --- |
| CHEMBL267099 |  | NS5 (Zika) | HPIV-3 | Zika and other flaviviruses (cell-based assays)<br>NS5 homologs in other flaviviruses (target-based) | Cell-based assays against Zika and other flaviviruses to confirm NS5 inhibition as a possible mechanism of action<br>Active against NS5 of Zika that could be tested against NS5 homologs from other flaviviruses |
|  |  |  |  | A panel of HPIV-3 targets | Explore possible mechanism of action for HPIV-3 inhibition observed in a cell-based assay |
| CHEMBL4437334 |  | NS2B-NS3 Protease (Dengue, Zika) | Dengue Virus, West Nile Virus, Yellow Fever Virus, Zika Virus | NS2B-NS3 homologs from other flaviviruses<br>(West Nile, Yellow Fever) | Cell-based and target-based assays support the hypothesis of activity against flaviviruses by targeting NS2B-NS3 |
| CHEMBL4522006 |  | NS2B-NS3 Protease (Dengue) | Dengue Virus | NS2B-NS3 homologs from other flaviviruses<br>Other flaviviruses in cell-based assays | Cell-based and target-based assays support the hypothesis of activity against flaviviruses by targeting NS2B-NS3 |
| CHEMBL1324 |  | NS2B-NS3 Protease (Dengue) | Dengue Virus | NS2B-NS3 homologs from other flaviviruses<br>Other flaviviruses in cell-based assays | Cell-based and target-based assays support the hypothesis of activity against flaviviruses by targeting NS2B-NS3 |
| CHEMBL3741713 |  | NS2B-NS3<br>Protease<br>(Dengue) | Dengue<br>Virus | NS2B-NS3<br>homologs<br>from other<br>flaviviruses<br>Other<br>flaviviruses<br>in cell-based<br>assays | Cell-based<br>and target-based<br>assays<br>support the<br>hypothesis<br>of activity<br>against<br>flaviviruses<br>by<br>targeting<br>NS2B-NS3 |
| CHEMBL4440832 |  | NS2B-NS3<br>Protease<br>(Dengue,<br>Zika) | Dengue<br>Virus | NS2B-NS3<br>homologs<br>from other<br>flaviviruses<br>Other<br>flaviviruses<br>in cell-based<br>assays | Cell-based<br>and target-based<br>assays<br>support the<br>hypothesis<br>of activity<br>against<br>flaviviruses<br>by<br>targeting<br>NS2B-NS3 |
| CHEMBL4462325 |  | NS2B-NS3<br>Protease<br>(Zika) | Dengue<br>Virus | NS2B-NS3<br>homologs<br>from other<br>flaviviruses<br>Other<br>flaviviruses<br>in cell-based<br>assays | Cell-based<br>and target-based<br>assays<br>support the<br>hypothesis<br>of activity<br>against<br>flaviviruses<br>by<br>targeting<br>NS2B-NS3 |
| CHEMBL4583315 |  | NS2B-NS3<br>Protease<br>(Zika) | Dengue<br>Virus | NS2B-NS3<br>homologs<br>from other<br>flaviviruses<br>Other<br>flaviviruses<br>in cell-based<br>assays | Cell-based<br>and target-based<br>assays<br>support the<br>hypothesis<br>of activity<br>against<br>flaviviruses<br>by<br>targeting<br>NS2B-NS3 |
| CHEMBL1370977 |  | NS2B-NS3<br>Protease<br>(Dengue) | Dengue<br>Virus | NS2B-NS3<br>homologs<br>from other<br>flaviviruses<br>Other<br>flaviviruses<br>in cell-based<br>assays | Cell-based<br>and target-based<br>assays<br>support the<br>hypothesis<br>of activity<br>against<br>flaviviruses<br>by<br>targeting<br>NS2B-NS3 |
| CHEMBL1458891 |  | NS2B-NS3<br>Protease<br>(Dengue) | Dengue<br>Virus | NS2B-NS3<br>homologs<br>from other<br>flaviviruses<br>Other<br>flaviviruses<br>in cell-based<br>assays | Cell-based<br>and target-based<br>assays<br>support the<br>hypothesis<br>of activity<br>against<br>flaviviruses<br>by<br>targeting<br>NS2B-NS3 |
| CHEMBL1980535 |  | NS2B-NS3<br>Protease<br>(Dengue) | Dengue<br>Virus | NS2B-NS3<br>homologs<br>from other<br>flaviviruses<br>Other<br>flaviviruses<br>in cell-based<br>assays | Cell-based<br>and target-based<br>assays<br>support the<br>hypothesis<br>of activity<br>against<br>flaviviruses<br>by<br>targeting<br>NS2B-NS3 |
| CHEMBL3628278 |  | NS2B-NS3<br>Protease<br>(West<br>Nile) | Dengue<br>Virus | NS2B-NS3<br>homologs<br>from other<br>flaviviruses<br>Other<br>flaviviruses<br>in cell-based<br>assays | Cell-based<br>and target-based<br>assays<br>support the<br>hypothesis<br>of activity<br>against<br>flaviviruses<br>by<br>targeting<br>NS2B-NS3 |
| CHEMBL3741422 |  | NS2B-NS3<br>Protease<br>(Dengue,<br>West Nile) | Dengue<br>Virus | NS2B-NS3<br>homologs<br>from other<br>flaviviruses<br>Other<br>flaviviruses<br>in cell-based<br>assays | Cell-based<br>and target-based<br>assays<br>support the<br>hypothesis<br>of activity<br>against<br>flaviviruses<br>by<br>targeting<br>NS2B-NS3 |
| CHEMBL4446364 |  | NS2B-NS3<br>Protease<br>(Zika) | Zika<br>Virus | NS2B-NS3<br>homologs<br>from other<br>flaviviruses<br>Other<br>flaviviruses<br>in cell-based<br>assays | Cell-based<br>and target-based<br>assays<br>support the<br>hypothesis<br>of activity<br>against<br>flaviviruses<br>by<br>targeting<br>NS2B-NS3 |
| CHEMBL4447165 |  | NS2B-NS3<br>Protease<br>(Zika) | Zika<br>Virus | NS2B-NS3<br>homologs<br>from other<br>flaviviruses<br>Other<br>flaviviruses<br>in cell-based<br>assays | Cell-based<br>and target-based<br>assays<br>support the<br>hypothesis<br>of activity<br>against<br>flaviviruses<br>by<br>targeting<br>NS2B-NS3 |
| CHEMBL4447800 |  | NS2B-NS3<br>Protease<br>(Dengue) | Dengue<br>Virus | NS2B-NS3<br>homologs<br>from other<br>flaviviruses<br>Other<br>flaviviruses<br>in cell-based<br>assays | Cell-based<br>and target-based<br>assays<br>support the<br>hypothesis<br>of activity<br>against<br>flaviviruses<br>by<br>targeting<br>NS2B-NS3 |
| CHEMBL4448497 |  | NS2B-NS3<br>Protease<br>(Zika) | Zika<br>Virus | NS2B-NS3<br>homologs<br>from other<br>flaviviruses<br>Other<br>flaviviruses<br>in cell-based<br>assays | Cell-based<br>and target-based<br>assays<br>support the<br>hypothesis<br>of activity<br>against<br>flaviviruses<br>by<br>targeting<br>NS2B-NS3 |
| CHEMBL4457232 |  | NS2B-NS3<br>Protease<br>(Zika) | Zika<br>Virus | NS2B-NS3<br>homologs<br>from other<br>flaviviruses<br>Other<br>flaviviruses<br>in cell-based<br>assays | Cell-based<br>and target-based<br>assays<br>support the<br>hypothesis<br>of activity<br>against<br>flaviviruses<br>by<br>targeting<br>NS2B-NS3 |
| CHEMBL4532866 |  | NS2B-NS3<br>Protease<br>(Zika) | Zika<br>Virus | NS2B-NS3<br>homologs<br>from other<br>flaviviruses<br>Other<br>flaviviruses<br>in cell-based<br>assays | Cell-based<br>and target-based<br>assays<br>support the<br>hypothesis<br>of activity<br>against<br>flaviviruses<br>by<br>targeting<br>NS2B-NS3 |
| CHEMBL4576745 |  | NS2B-NS3<br>Protease<br>(Dengue,<br>West Nile) | Dengue<br>Virus | NS2B-NS3<br>homologs<br>from other<br>flaviviruses<br>Other<br>flaviviruses<br>in cell-based<br>assays | Cell-based<br>and target-based<br>assays<br>support the<br>hypothesis<br>of activity<br>against<br>flaviviruses<br>by<br>targeting<br>NS2B-NS3 |
| CHEMBL522355 |  | NS2B-NS3<br>Protease<br>(West Nile) | West Nile<br>Virus | NS2B-NS3<br>homologs<br>from other<br>flaviviruses<br>Other<br>flaviviruses<br>in cell-based<br>assays | Cell-based<br>and target-based<br>assays<br>support the<br>hypothesis<br>of activity<br>against<br>flaviviruses<br>by<br>targeting<br>NS2B-NS3 |
| CHEMBL4454990 |  | NS5 (Zika) | Dengue<br>Virus,<br>Zika<br>Virus | NS5<br>homologs<br>from other<br>flaviviruses<br>Other<br>flaviviruses<br>in cell-based<br>assays | Cell-based<br>and target-based<br>assays<br>support the<br>hypothesis<br>of activity<br>against<br>flaviviruses<br>by<br>targeting<br>NS5 |
| CHEMBL4439416 |  | NS5 (Zika) | Zika<br>Virus | NS5<br>homologs<br>from other<br>flaviviruses<br>Other<br>flaviviruses<br>in cell-based<br>assays | Cell-based<br>and target-based<br>assays<br>support the<br>hypothesis<br>of activity<br>against<br>flaviviruses<br>by<br>targeting<br>NS5 |
| CHEMBL82242 |  | NS5<br>(Dengue) | Dengue<br>Virus | NS5<br>homologs<br>from other<br>flaviviruses<br>Other<br>flaviviruses<br>in cell-based<br>assays | Cell-based<br>and target-based<br>assays<br>support the<br>hypothesis<br>of activity<br>against<br>flaviviruses<br>by<br>targeting<br>NS5 |
| CHEMBL269277 |  | NS5<br>(Dengue) | Dengue<br>Virus | NS5<br>homologs<br>from other<br>flaviviruses | Cell-based<br>and target-based<br>assays<br>support the<br>hypothesis |
|  |  |  |  | Other flaviviruses in cell-based assays | of activity against flaviviruses by targeting NS5 |
| CHEMBL4449109 |  | NS5 (Dengue) | Dengue Virus | NS5 homologs from other flaviviruses<br>Other flaviviruses in cell-based assays | Cell-based and target-based assays support the hypothesis of activity against flaviviruses by targeting NS5 |
| CHEMBL4526128 |  | NS5 (Dengue) | Dengue Virus | NS5 homologs from other flaviviruses<br>Other flaviviruses in cell-based assays | Cell-based and target-based assays support the hypothesis of activity against flaviviruses by targeting NS5 |
| CHEMBL4587069 |  | NS5 (Dengue) | Dengue Virus | NS5 homologs from other flaviviruses<br>Other flaviviruses in cell-based assays | Cell-based and target-based assays support the hypothesis of activity against flaviviruses by targeting NS5 |

In other cases, as we observed for CHEMBL4437334, a compound was active against several viruses of the same family in phenotypic assays (Dengue Virus, West Nile Virus, Yellow Fever Virus, Zika Virus) and only tested and active against a subset of those viruses in the target-based assays (NS2B-NS3 Protease of Dengue and Zika). In these cases, we could suggest the compound be tested against the same target in the untested yet highly homologous viruses from the same family, using the principle of viral protein conservation.^3^

There were also instances of compounds, such as CHEMBL267099 (tubercidin), reported active in a phenotypic assay for a virus in one family (HPIV-3) and active in a target-based assay of another (Zika NS5 protein). Cases like these are particularly interesting, because after making this connection one can suggest testing this compound in various HPIV-3 targets, live cell Zika virus, as well as the other highly homologous *Flaviviridae* members (West Nile, Yellow Fever, Dengue) in live cell assay and against the NS5 protein.

Our concordance analysis also revealed 52 compounds active in at least one phenotypic assay and inactive in a (supposedly, relevant) target-based assay (**Table S5**, Supplementary Material). In this case, we recommend compounds be tested in additional target-based assays of the same viral family; it is also possible that the activity of such compounds inactive in viral targeting assays but active in phenotypic assays is actually due to their host-directed mechanism of action. There were also 191 compounds inactive in a phenotypic assay and active in at least one target-based assay (**Table S6**, Supplementary Material). Mapping the virus and viral family of the phenotypic result to the active result of the target helped identify potential new viral families for phenotypic testing, as well as highlighted the importance whether the cell type used in the phenotypic assay appropriately represented the virus and the antiviral result. Of course, due to the proportion of inactive compounds in our dataset, most compounds (5,656) were concordantly reported as inactive in both phenotypic and target-based assays.

### Integrated, searchable SMACC Database

We have described above various protocols for curating data of interest to the antiviral drug discovery from ChEMBL database. The resulting SMACC database currently exists as a searchable Excel spreadsheet that we include for public with this paper. This spreadsheet allows multiple approaches to identify compounds of interest. The approach described here in Methods, i.e., removing compounds with conflicting activity calls from our final BSA activity analysis, was stringent and resulted in a concise list of potential BSA compounds that we had the highest confidence in. While the filtering criteria described above was appropriate to achieve our project’s objective, we acknowledge the value of extracting different subsets of the SMACC database using different criteria depending on the study objectives, and SMACC database (even in the form of an Excel spreadsheet) enables multiple analyses.

For instance, in contrast with the approach described above, another method for identifying BSA compounds would be to consider all compounds with at least one active assay result against two or more viruses. This would increase the number of compounds considered in the analysis because inconclusive entries would not be removed as described above. To do this, we created a subset of the SMACC database with all compound entries reported as active in phenotypic assays, enumerated the number of viruses the compounds were reported active against, and removed all compounds reported active against only one virus. This approach resulted in 21 new hit compounds identified from phenotypic assays (**Table S7**) and 10 new hit compounds from target-target based assays that were not identified in the previous approach (**Table S8**).

As mentioned above, there are currently only 90 antiviral drugs approved for treating nine human infectious diseases.^4^ We utilized SMACCs filtering tools to enumerate their presence in our database. Currently, SMACC includes chemogenomics data for RSV and two strains of human influenza virus (H1N2 and H7N7), which covers only two of nine diseases with approved drugs. Despite this, we identified 53 of 90 approved drugs in our phenotypic dataset and 57 of 90 in our target-based dataset (**Table S9**). The compounds with reported active assay results are summarized in **Table 6**. Clearly, these drugs have broader activity than they are approved for. Further experimental testing based on hypotheses from this table will be extremely valuable to understanding their broad-spectrum potential.

**Table 6.** Approved drugs with active assay results found in SMACC.

| Compound Information |  |  |  |  | Active Assay Results in SMACC |  |
| --- | --- | --- | --- | --- | --- | --- |
| Drug name | Brand name | Approved clinical use | Inhibitory MOA | ChEMBL ID | Phenotypic Assay | Target-Based Assay |
| Simeprevir | Olysio® | HCV | NS3/NS4B Protease | CHEMBL501849 |  | H7N7 Matrix Protein 2 |
| Asunaprevir | Sunvepra® | HCV | Protease | CHEMBL2105735 |  | H7N7 Matrix Protein 2 |
| Sofosbuvir | Sovaldi® | HCV | NS5B | CHEMBL1259059 | Dengue Virus |  |
| Ribavirin | Copegus® | HCV, RSV, fever | RdRp | CHEMBL1643 | HPIV-3, Sandfly Fever, Dengue Virus, Yellow Fever Virus, RSV |  |
| Lopinavir | Kaletra® | HIV | Protease | CHEMBL729 | SARS-CoV-2 |  |
| Nelfinavir | Viracept® | HIV | Protease | CHEMBL584 | Dengue Virus |  |
| Raltegravir | Isentress® | HIV | Integrase | CHEMBL254316 |  | H7N7 Matrix Protein 2 |
| Elvitegravir | Vitekta® | HIV | Integrase | CHEMBL204656 |  | SARS-CoV-2 3CLpro |
| Atazanavir | Reyataz® | HIV | Protease | CHEMBL1163 |  | SARS-CoV-2 3CLpro |
| Rilpivirine | Edurant® | HIV-1 | Nonnucleoside reverse transcriptase | CHEMBL175691 | SARS-CoV-2 |  |
| Podofilox | Condylox® | HPV-related diseases | Cytotoxicity/cell division | CHEMBL61 | SARS-CoV-2 |  |
| Trifluridine | Viroptic® | HSV | Viral and cellular DNA synthesis | CHEMBL1129 | HPIV-3 |  |
| Idoxuridine | Dendrid® | HSV-1 | Viral and cellular DNA synthesis | CHEMBL788 |  | Zika NS5 |
| Acyclovir | Zovirax® | HSV, VZV | Viral DNA polymerase | CHEMBL184 |  | H7N7 Neuraminidase |
| Zanamivir | Relenza® | Influenza A and B | Neuraminidase | CHEMBL222813 | H7N7 |  |

Our pilot-SMACC database is currently available at https://smacc.mml.unc.edu. Users will find freely downloadable excel sheets containing our phenotypic, target-based, and overlapping datasets including tabs containing subsets of active compounds selected from the approach described in Methods. These excel sheets were designed so that users could easily extract subsets of the database using various filtering options. These filters include molecule (ChEMBL ID, smiles, InChiKey), virus, cell or target type, activity (activity call, raw assay result), and assay type. We acknowledge the widely varied objectives across antiviral research and emphasize the versatility of this database.

## Discussion

We have collected, curated, and integrated all the chemogenomic data available for a subset of viruses of interest in ChEMBL to identify BSA compounds. We created a pilot version of the SMACC database based on ChEMBL data. This initial data collection and curation effort can guide future data collection to increase the clarity and accessibility of relevant information to a broader scientific community include additional data on other emerging viruses. SMACC database adds to a variety of other important datasets and databases such as SARS-CoV-2 Data Resource by PubMed (https://www.ncbi.nlm.nih.gov/sars-cov-2/), a comprehensive COVID-19 Data Portal by European Bioinformatics Institute (EBI) (https://www.covid19dataportal.org), a collection of over 20,000 screening results for compounds tested against SARS-CoV-2 in a special release of the ChEMBL database (https://www.ebi.ac.uk/chembl/), a portal of target specific and phenotypic screening results of chemical libraries in SARS-CoV-2 established by NCATS at the NIH^26^ (https://opendata.ncats.nih.gov/covid19/index.html), and COVID-specific tools and collections like CORD-19 (https://www.kaggle.com/allen-institute-for-ai/CORD-19-research-challenge), COKE,^27^ and COVID-KOP.^28^ These collections along with several research initiatives such as the Antiviral Program for Pandemics (https://www.niaid.nih.gov/research/antivirals) and the Rapidly Emerging Antiviral Drug Development Initiative (READDI; https://www.readdi.org) pushed the scientific community to work in an ‘open science’ format.

Efforts similar to ours have been made to collect antiviral data prior to the SARS-CoV-2 outbreak. There is a collection of antiviral activity data from ChEMBL with enhanced taxonomy annotations as a tool for studying the antiviral chemical space that the authors dubbed “Viral ChEMBL”.^2^ Viral ChEMBL was compiled using information collected on compounds related to many virus types (human, animal, plant) and additional curation was performed to the data by mapping lists for assay and target organism data and using a dictionary of virus-related terms. While this collection is quite valuable to the field, it is based on an old version of ChEMBL20 (released 2015, current release is ChEMBL29) and some data are not relevant to human disease. Thus, our database was collected and manually curated to provide a structured, annotated repository of all data available in the most current version of ChEMBL for viruses that hold the greatest risk for human contraction and pandemic potential.

The SMACC database also has the potential to guide more informed drug repurposing efforts, which was a popular strategy employed during the first year of the SARS-CoV-2 pandemic. Repurposing FDA approved drugs ^29^ and their combinations ^30^ quickly provided options for clinical use without the need to undergo extensive toxicological testing. For instance, we have identified anticancer drug brequinar as a potential antiviral agent (cf. Table 3), and the analysis of additional bioactivities, including those against host targets, may reveal novel interesting compounds.

Beyond drug repurposing, another intuitive approach used in the most recent SARS-CoV-2 pandemic to identify BSA drugs across the coronaviruses was through identifying proteome conservation, which was studied by Schapira et al.^31^ as well as our group.^3^ Schapira et al. analyzed the conservation of all available PDB structures of α- and β-coronaviruses, as well as samples from patients with SARS-CoV-2 by mapping druggable binding pockets onto experimental structures of SARS-CoV-2 proteins. Our work complemented that of Schapira et al.,^31^ by exploring the idea that similarities between homologous coronaviral proteins could be exploited for target selection and the development of broad-spectrum anti-coronaviral compounds. Putting that idea into the context of potential broad-spectrum inhibitors of conserved targets, we identified drugs from existing literature that inhibit M^pro^, RdRp, PL^pro^, and nsp10-nsp16, carefully collected and analyzed all known experimental data on their antiviral activity and validated our hypothesis by estimating their potential as broad-spectrum drugs. These compounds are discussed extensively elsewhere and are naturally included in the SMACC database. Thus, we feel exploring the conservation between homologous coronaviral proteins is an extremely valuable strategy for target selection and could assist the development of BSA compounds.

With viral protein conservation as a tool for identifying BSAs, one wonders if there may be a link between protein conservation and ligand promiscuity. While in theory the framework of our database would easily allow for this analysis, the unfortunate truth is that the data are not available, as our collection of target-based data was already far more limited than our phenotypic set. Further, it is no secret that merely collecting such data from available data sources can be misleading. Errors described above depict the challenges we faced in curation and collection; for example, a user looking for compounds active against NS2B-NS2 Protease would not have found results due to the target being annotated generally as “genome polyprotein.” We hope that our systematic analysis and enumeration of annotation deficiencies and bioactivity data curation protocols could help other researchers interested in expanding our collection or creating their own specialized collections. Most importantly, our efforts both identified several BSAs discovered by chance without deliberate focused efforts (cf. Tables 3-4) as well as nominated several compounds for additional testing (cf. Table 5). As discussed above, the SMACC database included with this paper, enables user-defined filtering of the data to support the generation of specialized subsets. In summary, we posit that this study provides strong motivation for continued investments into research targeting the discovery and development of novel BSA agents.

## Conclusions

We have developed a pilot version of the SMACC (Small Molecule Antiviral Compound Collection) database containing over 32,500 entries for 13 emerging viruses. We followed the following steps to create SMACC: (i) identification, collection, and curation of all chemical bioactivity data available in ChEMBL for 13 emerging viruses holding the greatest potential threat to global human health; (ii) identification and resolution of the data availability, reproducibility and quality challenges; (iii) integration of curated and carefully annotated data on compounds tested in both phenotypic (21,392 entries) and target-based (11,123 entries) assays for these viruses; and (iv) identification of chemicals showing high potential for BSA activity. Specifically, we identified eight compounds active against 3-4 viruses from the phenotypic data, 16 compounds active against two viruses from the target-based data, and 35 compounds active in at least one phenotypic and one target-based assay. Duplicates (phenotypic and overlap sets) and singletons (all sets) were also identified and annotated. While the pilot version of SMACC has integrated all chemogenomic data available in ChEMBL for these viruses, there was a large degree of sparsity (93%) within the integrated data matrix. Many viruses were understudied and thus, important results may be obtained by targeted testing of compounds included in SMACC against targets other than those against which they were tested. In fact, we have suggested several such targeted testing experiments in this paper (cf. Table 5).

Our analysis indicates that not many BSAs have emerged from previous disconcerted studies and that special, focused efforts must be established going forward. The SMACC database built in this study may serve as a reference for virologists and medicinal chemists working on the development of BSA agents in preparation for future viral outbreaks. The SMACC database is publicly available in the form of searchable Excel spreadsheet at https://smacc.mml.unc.edu.

## Acknowledgement

Authors from UNC-Chapel Hill were supported by National Institutes of Health (Grants U19AI171292 and R01GM140154). This research was supported by the Intramural Research Program of the National Center for Advancing Translational Sciences (NCATS), National Institutes of Health (NIH).

